# Defining Multiple Layers of Intratumor Heterogeneity Based on Variations of Perturbations in Multi-omics Profiling

**DOI:** 10.1101/2022.10.15.512357

**Authors:** Hongjing Ai, Dandan Song, Xiaosheng Wang

## Abstract

Intratumor heterogeneity (ITH) is associated with tumor progression, relapse, immunoevasion, and drug resistance. Existing algorithms for measuring ITH are limited to at a single molecular level. We proposed a set of algorithms for measuring ITH at the genome (somatic copy number alterations (CNAs) and mutations), mRNA, microRNA (miRNA), long non-coding RNA (lncRNA), protein, and epigenome level, respectively. These algorithms were designed based on a common concept: information entropy. By analyzing 33 TCGA cancer types, we demonstrated that these ITH measures had the typical properties of ITH, namely their significant correlations with unfavorable prognosis, tumor progression, genomic instability, antitumor immunosuppression, and drug resistance. Furthermore, we showed that the correlations between ITH measures at identical molecular levels were stronger than those at different molecular levels. The mRNA ITH showed stronger correlations with the miRNA, lncRNA, and epigenome ITH than with the genome ITH, supporting the regulatory relationships of miRNA, lncRNA, and DNA methylation towards mRNA. The protein ITH displayed stronger correlations with the transcriptome-level ITH than with the genome-level ITH, supporting the central dogma of molecular biology. Finally, we integrated the seven ITH measures into an ITH measure, which displayed more prominent properties of ITH than the ITH measures at a single molecular level. This analysis of multi-level ITH provides novel insights into tumor biology and potential values in clinical practice for pan-cancer.

## Introduction

Intratumor heterogeneity (ITH) is the variation of phenotypic and molecular profiles among different tumor cells within a tumor. Abundant evidence has shown that ITH is associated with tumor progression, immunoevasion, therapy resistance, and unfavorable outcomes [1]. Hence, quantitative analysis of ITH is advisable in the management of cancer patients. To this end, a number of algorithms have been developed to quantify ITH [2]. Among them, the algorithms for quantifying ITH at the genetic level are the most common, which measure ITH usually based on alterations in profiles of somatic mutations and/or copy number alterations (CNAs). Such algorithms include MATH [3], EXPANDS [4], PhyloWGS [5], and DITHER [6], to name a few. Nevertheless, although genetic or DNA-level ITH is common in cancer, it cannot fully reflect the phenotypic ITH [2]. Therefore, a few algorithms for quantifying ITH at the mRNA level have been proposed, such as DEPTH [2] and tITH [7]. In addition, an algorithm for quantifying ITH based on alterations in DNA methylation profiles has been designed [8]. These non-genetic ITH evaluation algorithms have been demonstrated to have comparable performance in characterizing the typical properties of ITH [2, 7, 8].

Information entropy (IE), a concept introduced by Shannon [9], representing disorder, uncertainty, or variation of variables, has been applied in a broad spectrum of subjects, including biology [10]. Since ITH represents the variation of phenotypic and molecular profiles among tumor cells, applying IE for characterizing ITH is a viable approach [6, 7]. For example, the DITHER algorithm measures ITH based on the IE of somatic mutation and CNA profiles in tumor cells [6]. tITH evaluates ITH by the IE of protein-protein interaction network [7]. These IE-based algorithms have been shown to be competitive in describing ITH as compared to other algorithms [6, 7].

In this study, we proposed a set of IE-based algorithms to quantify ITH based on profiles of somatic mutations, CNAs, DNA methylation, mRNA expression, microRNA (miRNA) expression, long non-coding RNA (lncRNA) expression, and protein expression, respectively. We tested these algorithms in 33 cancer types from The Cancer Genome Atlas (TCGA) program [11]. First, we analyzed correlations between the ITH scores and tumor clinical, phenotypic, and molecular features, including survival prognosis, tumor stage, grade, and metastasis, stemness, proliferation, antitumor immune response, and genomic instability. Second, we investigated pairwise correlations between the ITH scores by the different methods. Finally, we integrated all the ITH scores by the different methods into an ITH score and explored its association with tumor clinical, phenotypic, and molecular features. This study evaluated ITH based on multiple layers of molecular profiles and would provide novel insights into the biology of ITH as well as valuable clinical implications for the cancer management.

## Materials and Methods

### Materials

We obtained multi-omics and clinical data for 33 TCGA cancer types from the Genomic Data Commons (GDC) data portal (https://portal.gdc.cancer.gov/). The 33 cancer types included adrenocortical carcinoma (ACC), bladder urothelial carcinoma (BLCA), breast invasive carcinoma (BRCA), cervical squamous cell carcinoma and endocervical adenocarcinoma (CESC), cholangiocarcinoma (CHOL), colon adenocarcinoma (COAD), lymphoid neoplasm diffuse large B-cell lymphoma (DLBC), esophageal carcinoma (ESCA), glioblastoma multiforme (GBM), head and neck squamous cell carcinoma (HNSC), kidney chromophobe (KICH), kidney renal clear cell carcinoma (KIRC), kidney renal papillary cell carcinoma (KIRP), acute myeloid leukemia (LAML), brain lower grade glioma (LGG), liver hepatocellular carcinoma (LIHC), lung adenocarcinoma (LUAD), lung squamous cell carcinoma (LUSC), mesothelioma (MESO), ovarian serous cystadenocarcinoma (OV), pancreatic adenocarcinoma (PAAD), pheochromocytoma and paraganglioma (PCPG), prostate adenocarcinoma (PRAD), rectum adenocarcinoma (READ), sarcoma (SARC), skin cutaneous melanoma (SKCM), stomach adenocarcinoma (STAD), testicular germ cell tumors (TGCT), thyroid carcinoma (THCA), thymoma (THYM), uterine corpus endometrial carcinoma (UCEC), uterine carcinosarcoma (UCS), and uveal melanoma (UVM). These multi-omics data were the profiles of somatic mutations, CNAs, DNA methylation, mRNA expression, microRNA (miRNA) expression, long non-coding RNA (lncRNA) expression, and protein expression. We also downloaded somatic mutations-associated “maf” files and CNAs-associated “SNP6” files from the GDC data portal. A description of these data is shown in Supplementary Table S1.

### The IE-based algorithms for evaluating ITH

For each tumor sample, we calculated its IE-based ITH scores as follows:

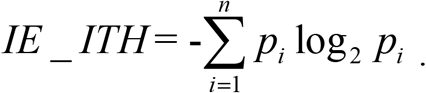

We calculated seven types of *IE* _ *ITH* based on profiles of somatic CNAs (cnvITH), somatic mutations (mutITH), DNA methylation (metITH), mRNA expression (mrnITH), miRNA (mirITH) expression, lncRNA (lncITH) expression, and protein expression (proITH), respectively. For cnvITH, *n* is the number of bins grouping mean segments (= log_2_(copy number (CN)/2)) in the tumor, and *p*_*i*_ is the proportion of mean segments in the *i*th bin [6]; for mutITH, *n* is the number of bins grouping the somatically mutant allele fractions (MAFs) among loci in the tumor, and *p*_*i*_ is the proportion of loci whose MAFs lie in the *i*th bin [6]; for metITH, *n* is the number of bins grouping DNA methylation probes, and *p*_*i*_ is the proportion of probes with the distances of β-values in the tumor from average β-values in normal controls (or in tumor samples if normal controls are not available) lie in the *i*th bin; for mrnITH, mirITH, lncITH, and proITH, *n* is the number of bins grouping RNAs or proteins, and *p*_*i*_ is the proportion of RNAs or proteins with the distances of their expression values in the tumor from average expression values in normal controls (or in tumor samples if normal controls are not available) lie in the *i*th bin. Consistently, we set *n* as 20 for the seven types of ITH evaluation algorithms. That is, mean segments, loci, methylation probes, mRNAs, miRNAs, lncRNAs, and proteins were grouped into 20 bins of equal width based on log_2_(CN/2), MAFs, or distances of expression values, where the *k*th bin is [*min* + (*k*-1) × (*max*-*min*)/20, *min* + *k* × (*max*-*min*)/20], where *min* and *max* denote the minimum and maximum log_2_(CN/2), MAFs, distances of β-values, or distances of expression values within the tumor. Before calculating ITH scores, we normalized distances of β-values and distances of expression values into the range [0, 1] by using the formula:

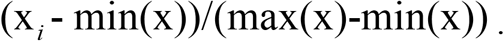

An illustration of the ITH algorithms is shown in Figure 1. Finally, for each tumor, we integrated its seven ITH measures into an ITH measure by taking the average of the seven ITH scores after scaling to the range [0, 1]. The R package for the IE-based ITH measure algorithms is available at the website: https://github.com/WangX-Lab/ieITH, under a GNU GPL open-source license.

**Fig. 1.**
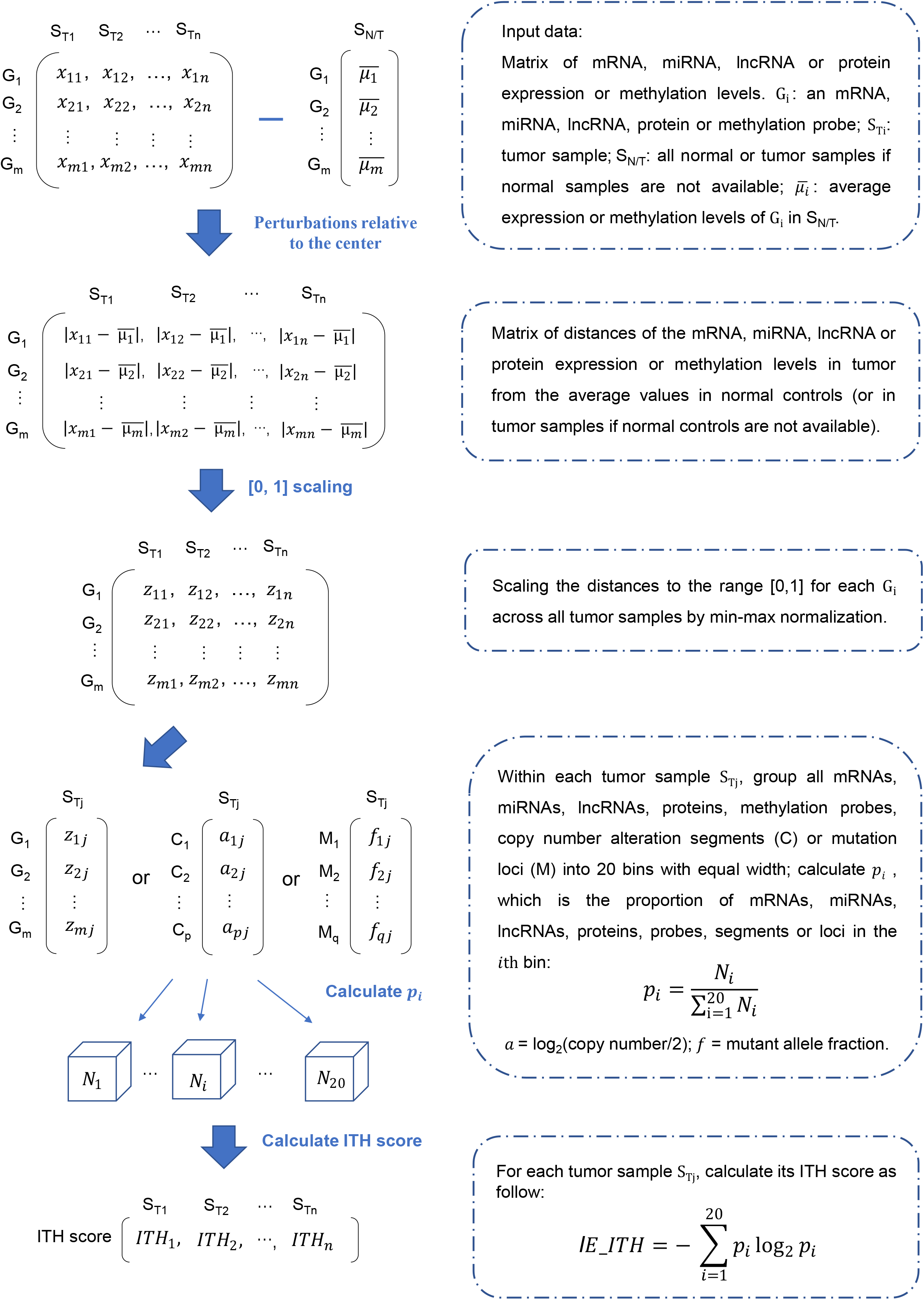
Illustration of the IE-based ITH algorithms.

### Gene-set enrichment analysis

We calculated the enrichment scores of immune signatures, biological processes (such as tumor stemness and cell proliferation), and pathways in a tumor by the single-sample gene-set enrichment analysis (ssGSEA) [12] based on the expression profiles of their marker genes or pathway genes. The ratio of two immune signatures in a tumor was defined as log2-transformed values of the geometric mean expression level of all marker genes in an immune signature divided by that in another immune signature. The marker genes or pathway genes of immune signatures, biological processes, and pathways are presented in Supplementary Table S2.

### Quantification of tumor mutation burden (TMB), homologous recombination deficiency (HRD), CNA, and tumor purity

For a tumor, its TMB was defined as its total count of somatic mutations; the CNA score, also known as ploidy score, was calculated by the ABSOLUTE algorithm [13] with the input of “SNP6” files; The HRD combined scores of loss of heterozygosity, large-scale state transitions and the number of telomeric allelic imbalances. We obtained the HRD scores of tumors from the publication by Knijnenburg et al [14]. Tumor purity was estimated by the ESTIMATE algorithm [15] with the input of gene expression profiles.

### Clustering analysis

We identified subgroups of pan-cancer based on the seven types of ITH scores by using the two-way hierarchical clustering method. Before clustering, we normalized the ITH scores by z-score and transformed them into distance matrices by the R function “dist” (the “method” parameter = “euclidean”). We implemented the hierarchical clustering with the R function “hclust” in the R package “stats” with the parameters: method = “ward.D2” and members = NULL. To quantify the distribution of tumor samples in each cancer across defined subgroups, we applied the chi-squared test to assess whether the distribution significantly deviates from random expectations. Further, we calculated the Ro/e for each combination of cancers and subgroups. The Ro/e was defined as the ratio of the observed sample size to the expected sample size of a given combination of cancer and subgroup. The largest Ro/e indicates that a cancer type is most enriched in a specific subgroup.

### Survival analysis

We compared survival prognosis between different classes of cancer patients by using the Kaplan-Meier (KM) model [16]. The survival prognosis involved four types of clinical endpoints: overall survival (OS), disease-specific survival (DSS), disease-free interval (DFI), and progression-free interval (PFI). We utilized KM curves to display survival time differences between high-ITH (ITH scores in the upper third) and low-ITH (ITH scores in the bottom third) cancer patients, or different groups of cancer patients clustered by the seven ITH scores. The log-rank test was employed to evaluate the significance of survival time differences. The R function “survfit” in the R package “survival” was utilized to perform this kind of survival analysis.

### ITH scores by other methods

To compare the IE-based ITH measures with other ITH measures, we also calculated ITH scores of the TCGA tumors by other methods, including MATH [3], PhyloWGS [5], DITHER [6], DEPTH [2], and MYTH [8]. We evaluated the ITH scores by MATH with the R function “math.score” in the R package “maftools” with the input of “maf” files. MATH defines a tumor’s ITH score as the ratio of the median absolute deviation to the median of its MAFs [3]. The ITH scores by PhyloWGS were obtained from the associated publication [5]. PhyloWGS evaluates ITH by inferring the subclones of tumor cells based on their somatic mutations and CNAs [5]. Likewise, DITHER calculates a tumor’s ITH score based on both its somatic mutation and CNA profiles [6]. We evaluated the ITH scores by DITHER using the R package “DITHER” with the input of “maf” and “SNP6” files. DEPTH evaluates ITH at the mRNA level [2]. We calculated DEPTH ITH scores with the R package “DEPTH” with the input of gene expression matrix. MYTH calculates the ITH index of tumors based on their DNA methylation profiles [8]. We calculated MYTH ITH scores using the R package “MYTH” with the input of gene-level DNA methylation values.

### Statistical analysis

We compared ITH scores between two groups of samples by the one-tailed Mann–Whitney *U* test. In evaluating correlations among ITH scores and between ITH scores and other variables, we utilized the Spearman method to report *p* values and correlation coefficients (*ρ*). In analyzing contingency tables, we employed the chi-square test. We performed all statistical analysis with the R programing language (version 4.1.1).

## Results

### The IE-based ITH likely correlates with unfavorable clinical outcomes and tumor progression phenotypes in cancer

We analyzed correlations between the seven types of ITH scores and survival prognosis in pan-cancer and in several common cancer types, including breast cancer (BRCA), lung cancer (LUAD and LUSC), gastrointestinal (GI) cancer (ESCA, STAD, COAD, and READ), kidney cancer (KIRC, KICH, and KIRP), melanoma (SKCM), and glioma (LGG). In pan-cancer, five types of ITH (cnvITH, mutITH, metITH, lncITH, and mrnITH) scores displayed significant negative correlations with OS, DSS, DFI, and PFI, and mirITH had significant negative correlations with DSS, DFI, and PFI (log-rank test, *p* < 0.05) (Fig. 2A). In addition, proITH had a negative correlation with DSS (*p* = 0.07) (Fig. 2A). In the common cancer types, the ITH scores were likely to correlated significantly and negatively with survival prognosis, particularly cnvITH, lncITH, and mrnITH (Supplementary Fig. S1A). For example, in LGG, six of the seven types of ITH (except mutITH) scores had significant negative correlations with OS, DSS, and PFI (Supplementary Fig. S1A). Overall, these results suggest that the IE-based ITH scores are likely to be adverse prognostic factors.

**Fig. 2.**
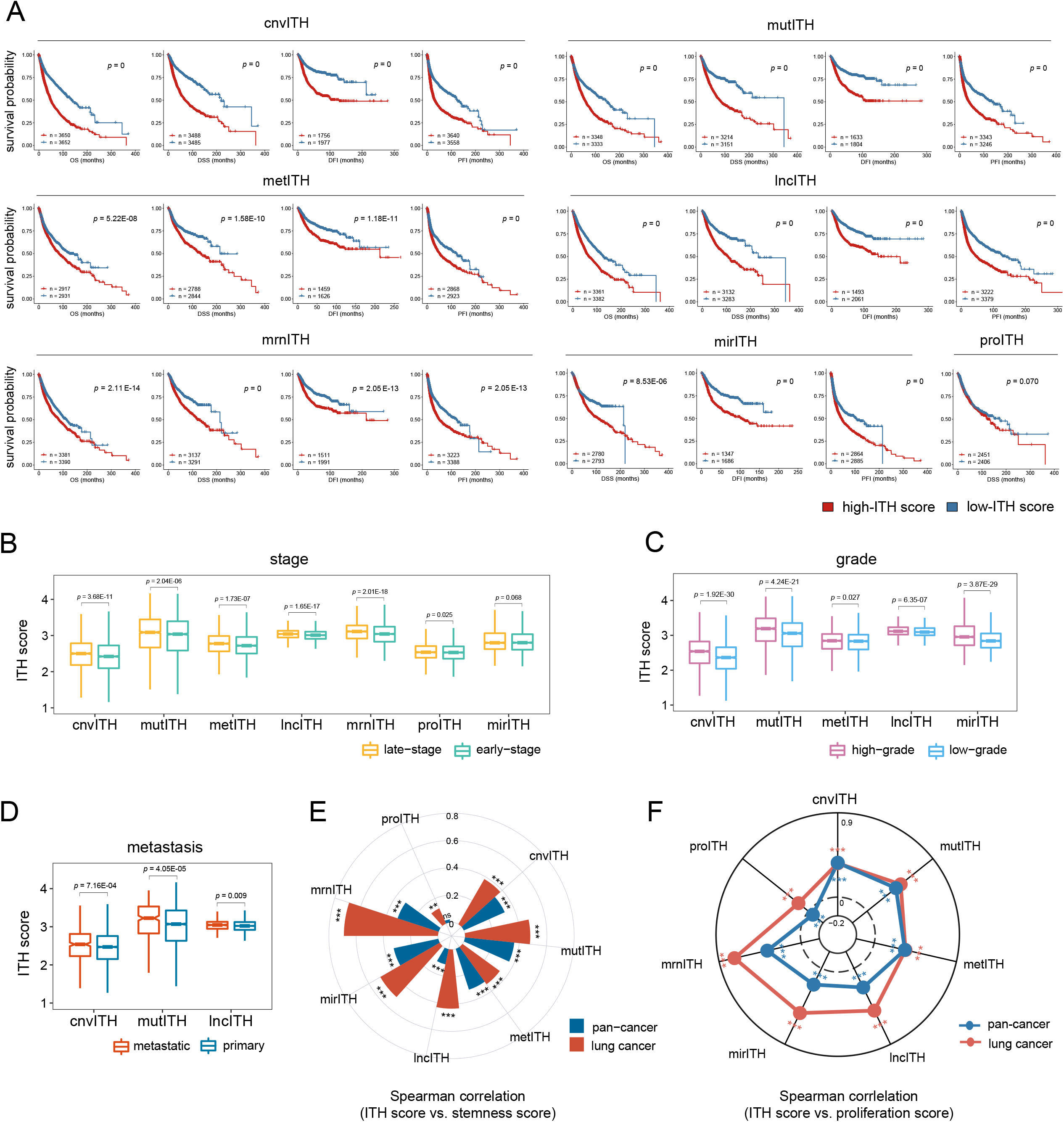
Correlations of the IE-based ITH measures with clinical and phenotypic features in cancer. **(A)** KM curves showing that patients with high-ITH scores (upper third) have better survival than those with low-ITH scores (bottom third) in pan-cancer. The log-rank test *p* values are shown. OS, overall survival. DSS, disease-specific survival. DFI, disease-free interval. PFI, progression-free interval. DFS, disease-free survival. **(B-D)** Seven types of ITH scores are significantly higher in late-stage (stage III-IV) than in early-stage (stage I-II) **(B)**, five types of ITH scores are significantly higher in high-grade (G3-4) versus low-grade (G1-2) **(C)**, and three types of ITH scores are significantly higher in metastatic versus primary tumors in pan-cancer **(D)**. The one-tailed Mann–Whitney *U* test *p* values are shown. **(E, F)** IE-based ITH scores are positively correlated with tumor stemness scores **(E)** and tumor cell proliferation scores **(F)** in pan-cancer and in lung cancer. The Spearman correlation coefficients are shown. * *p* < 0.05, ** *p* < 0.01, *** *p* < 0.001; they also apply to the following figures.

Tumor stage represents the stage of tumor development. Notably, six types of ITH (cnvITH, mutITH, metITH, lncITH, mrnITH, and proITH) scores were significantly higher in late-stage (stage III-IV) than in early-stage (stage I-II) tumors in pan-cancer (one-tailed Mann–Whitney *U* test, *p* < 0.05); mirITH scores were higher in late-stage than in early-stage tumors in pan-cancer (*p* = 0.068) (Fig. 2B). In individual cancer types, such as lung cancer and kidney cancer, most ITH scores were significantly higher in late-stage versus early-stage tumors (Supplementary Fig. S1B). Tumor grade reflects the abnormality degree of tumor cells versus normal cells and indicates how fast a tumor is likely to progress. We found that five types of ITH (cnvITH, mutITH, metITH, lncITH, and mirITH) scores were significantly higher in high-grade (G3-4) than in low-grade (G1-2) tumors in pan-cancer (*p* < 0.05) (Fig. 2C). Besides, in LGG and kidney cancer, there were 7 and 6 ITH scores were significant higher in high-grade than in low-grade tumors (*p* < 0.05) (Supplementary Fig. S1C). Furthermore, we found that three types of ITH (cnvITH, mutITH, and lncITH) scores were significantly higher in metastatic than in primary tumors in pan-cancer (*p* < 0.05) (Fig. 2D). In addition, four types of ITH scores were significant higher in high-grade than in low-grade tumors in kidney cancer (*p* < 0.01) (Supplementary Fig. S1D). Altogether, these results demonstrate that the IE-based ITH scores likely increase with tumor progression.

Tumor stemness refers to stem cell-like properties of cancer cells, which have essential associations with ITH, tumor progression, relapse, and therapy resistance [17]. We observed that six types of ITH (cnvITH, mutITH, metITH, lncITH, mirITH, and mrnITH) scores were positively correlated with tumor stemness scores in pan-cancer (Spearman correlation, *p* < 0.001) (Fig. 2E). In lung cancer, SKCM, LGG, and BRCA, there were 7, 7, 6, and 6 ITH scores showing significant positive correlations with tumor stemness scores, respectively (*p* < 0.05) (Fig. 2E and Supplementary Fig. S1E). Meanwhile, six types of ITH (cnvITH, mutITH, metITH, lncITH, mirITH, and mrnITH) scores had significant positive correlations with tumor cell proliferation scores in pan-cancer (*p* < 0.001) (Fig. 2F). In lung cancer, LGG, and BRCA, all the seven types of ITH scores were positively correlated with tumor cell proliferation scores (*p* < 0.02) (Fig. 2F and Supplementary Fig. S1F).

Altogether, these results suggest that the IE-based ITH is associated with unfavorable clinical outcomes in cancer.

### The IE-based ITH likely correlates with genomic instability in cancer

Genomic instability has been indicated as a major cause of ITH [18]. Tumor aneuploidy, also known as CNAs, often results from genomic instability [19]. We found that six ITH scores correlated positively with CNA scores in pan-cancer (Spearman correlation, *p* < 0.001) (Fig. 3A). Notably, cnvITH had the strongest correlation with CNA scores (*ρ* = 0.72), followed by mutITH (*ρ* = 0.60) and metITH (*ρ* = 0.36). It is justified since cnvITH, mutITH, and metITH are the ITH measure at the DNA level. Moreover, in the common cancer types, most of the ITH scores had significant positive correlations with CNA scores (Supplementary Fig. S2A). Increased tumor mutation loads are also a consequence of genomic instability [18]. We found that five ITH (cnvITH, mutITH, metITH, lncITH, and mrnITH) scores correlated positively with TMB in pan-cancer (*p* < 0.001) (Fig. 3B). Also, most of the ITH scores had significant positive correlations with TMB in the common cancer types (Supplementary Fig. S2A). HRD may lead to large-scale genomic instability [14]. We observed that six types of ITH (cnvITH, mutITH, metITH, lncITH, mirITH, and mrnITH) scores had significant positive correlations with HRD scores (*p* < 0.001) (Fig. 3C). In BRCA, lung cancer, GI cancer, and SKCM, there were 7, 7, 6, and 5 ITH scores showing significant positive correlations with HRD scores, respectively (*p* < 0.05) (Supplementary Fig. S2A).

**Fig. 3.**
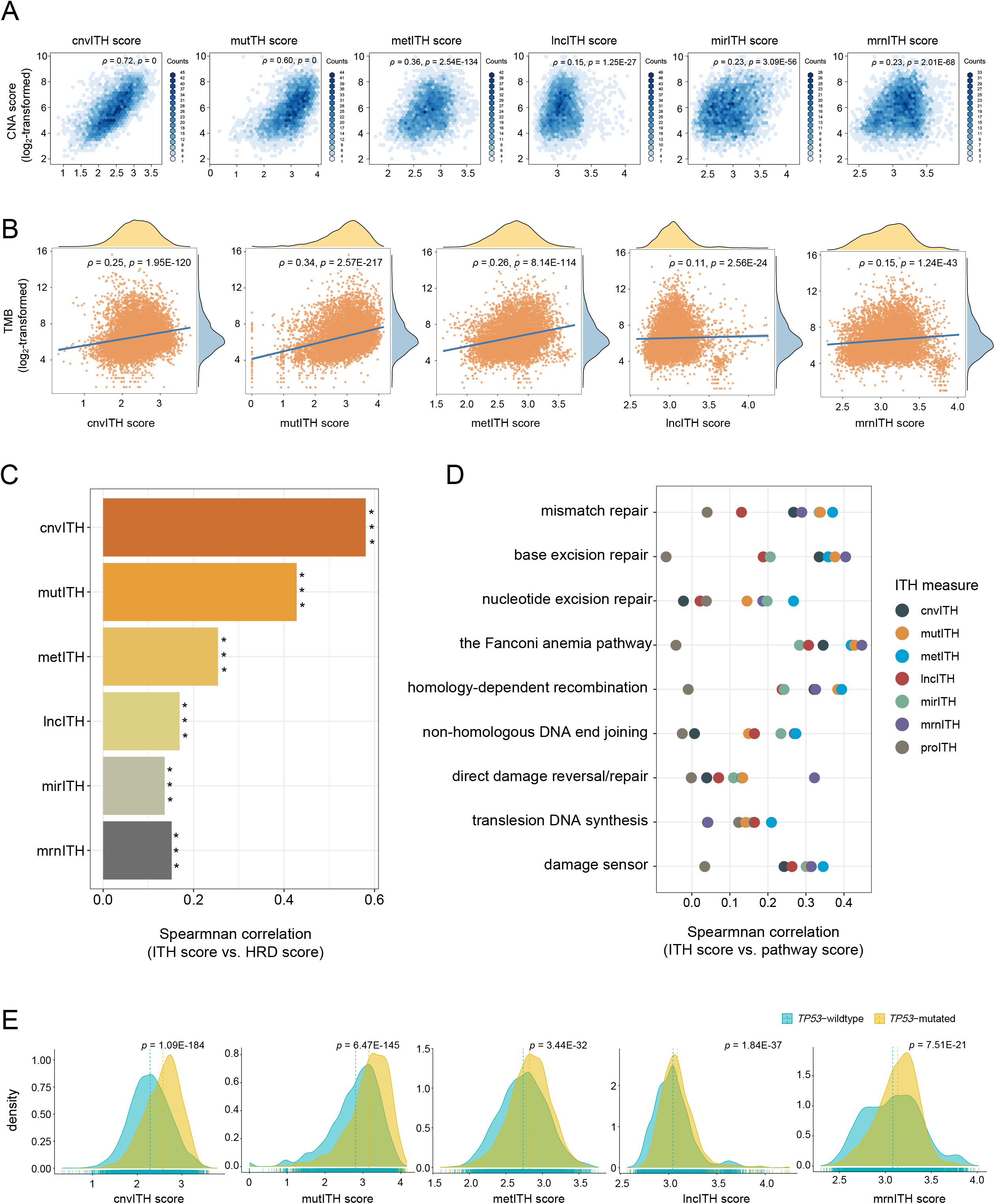
Correlations between the IE-based ITH scores and genomic instability. **(A)** Six types of ITH scores are positively correlated with CNA scores in pan-cancer. **(B)** Five types of ITH scores are positively correlated with TMB in pan-cancer. **(C)** Six types of ITH scores had positive correlations with HRD scores in pan-cancer. **(D)** ITH scores are positively correlated with the enrichment of DNA damage repair pathways in pan-cancer. The Spearman correlation coefficients and *p* values are shown in **(A-D). (E)** Five types of ITH scores are significantly higher in *TP53*-mutated than in *TP53*-wildtype tumors in pan-cancer. The one-tailed Mann– Whitney *U* test *p* values are shown. TMB, tumor mutation burden. HRD, homologous recombination deficiency.

DNA damage repair (DDR) pathways are important in maintaining genomic stability, whose upregulation in cancer indicates genomic instability. We analyzed correlations between ITH scores and the enrichment scores of nine DDR pathways in pan-cancer and in lung cancer. The nine DDR pathways included mismatch repair, base excision repair, nucleotide excision repair, the Fanconi anemia (FA) pathway, homology-dependent recombination, non-homologous DNA end joining, direct damage reversal/repair, translesion DNA synthesis, and damage sensor [14]. Most of the pairwise correlations between these pathways’ enrichment and ITH scores were significant and positive (*p* < 0.05) (Fig. 3D and Supplementary Fig. S2B). Notably, in pan-cancer, metITH scores had the strongest positive correlations with six of the nine DNA damage repair pathways’ enrichment in the seven ITH measures (*p* < 0.05) (Fig. 3D).

*TP53* encodes one of the most important tumor suppressor, whose mutations are frequent across cancer [20]. Since *TP53* plays key roles in maintaining genomic integrity [21], its mutations may enhance genomic instability. We observed that five ITH (cnvITH, mutITH, metITH, lncITH, and mrnITH) scores were far higher in *TP53*-mutated than in *TP53*-wildtype tumors in pan-cancer (one-tailed Mann–Whitney *U* test, *p* < 0.001) (Fig. 3E). Similar results were observed in BRCA, lung cancer, and GI cancer (*p* < 0.01) (Supplementary Fig. S2C).

Taken together, these results support that the IE-based ITH has markedly correlations with genomic instability in cancer.

### The IE-based ITH likely correlates with antitumor immunosuppression

ITH plays key roles in antitumor immune evasion [1]. We analyzed correlations between ITH scores and the enrichment of two antitumor immune signatures (CD8+ T cells and cytolytic activity) in pan-cancer and in the common cancer types. We found that the correlations between the seven ITH scores and both immune signatures’ enrichment scores were all significant and negative in pan-cancer (Spearman correlation, *p* < 0.001) (Fig. 4A). In the common cancer types, most of the correlations were also significant and negative (Supplementary Fig. S3). Furthermore, we analyzed correlations between ITH scores and the ratios of immune-stimulatory/immune-inhibitory signatures (CD8+/CD4+ regulatory T cells). Likewise, the correlations between the seven ITH scores and the ratios were significant and negative in pan-cancer (*p* < 0.001) (Fig. 4B). In addition, 7, 7, and 6 ITH scores had significant negative correlations with the ratios of CD8+/CD4+ regulatory T cells in BRCA, lung cancer, and SKCM, respectively (Supplementary Fig. S3). Taken together, these results support the inverse correlation between the IE-based ITH and antitumor immunity in cancer.

**Fig. 4.**
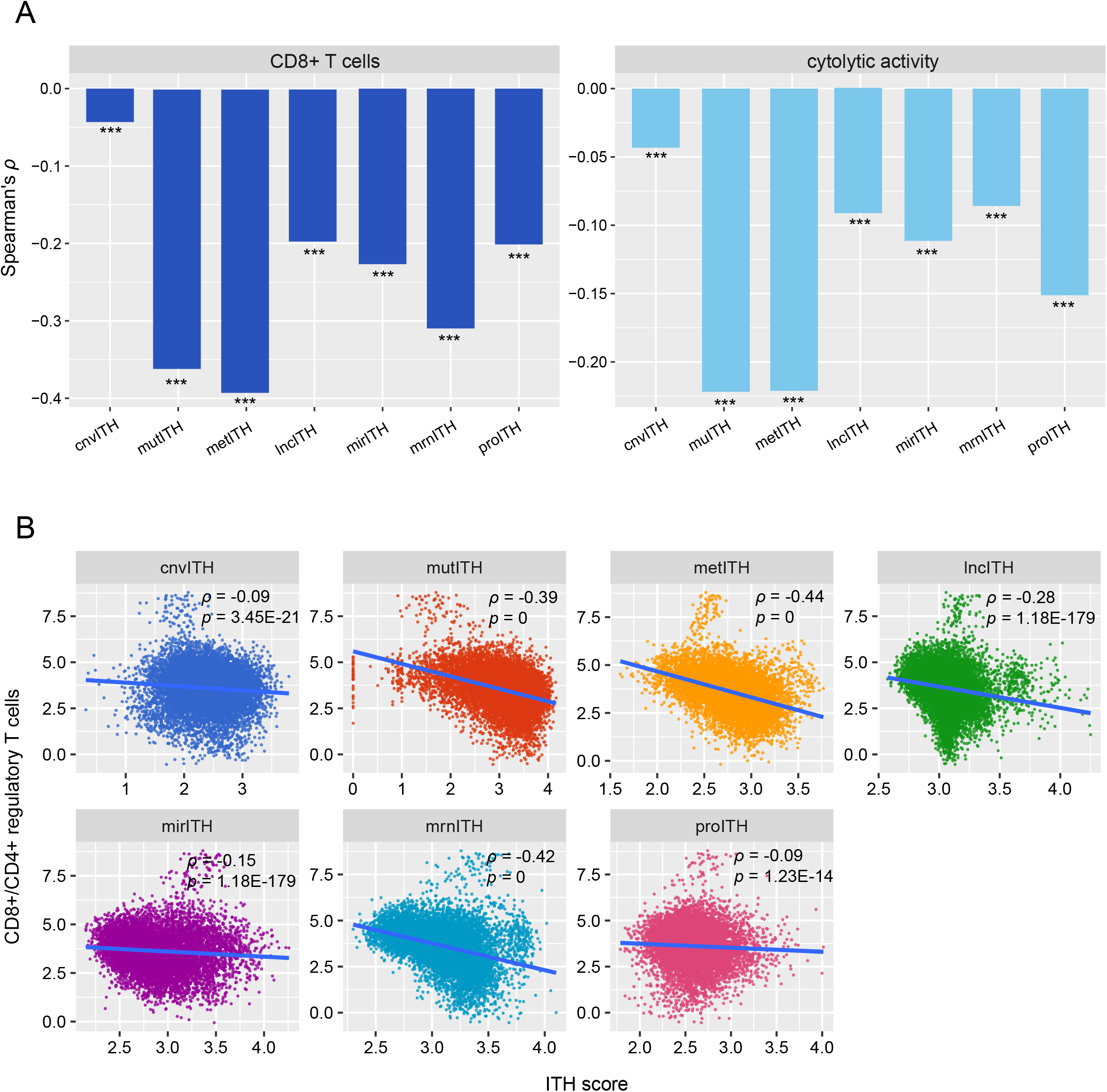
Correlations between the IE-based ITH scores and antitumor immune responses. The seven ITH scores are inversely correlated with the enrichment levels of CD8+ T cells and immune cytolytic activity **(A)**, and the ratios of CD8+/CD4+ regulatory T cells signatures **(B)** in pan-cancer. The Spearman correlation coefficients (*ρ*) and *p* values are shown.

### The IE-based ITH likely correlates with drug resistance

ITH may confer drug resistance in cancer therapies [22]. We analyzed the correlation between the IE-based ITH scores and drug responses in TCGA pan-cancer. If a patient had a complete or partial drug response, we classified him/her into the responsive group and otherwise non-responsive group. For the responses to chemotherapies or targeted therapies, six ITH scores (mutITH, metITH, lncITH, mirITH, mrnITH, and proITH) were significantly lower in the responsive than in the non-responsive group (*p* < 0.01) (Fig. 5A). For the response to immunotherapies, lncITH and proITH were significantly lower in the responsive than in the non-responsive group (*p* < 0.01) (Fig. 5B). Interestingly, cnvITH scores were by contrast significantly higher in the responsive than in the non-responsive group with respect to both chemotherapies or targeted therapies (*p* = 0.045) (Fig. 5A) and immunotherapies (*p* = 0.029) (Fig. 5B).

**Fig. 5.**
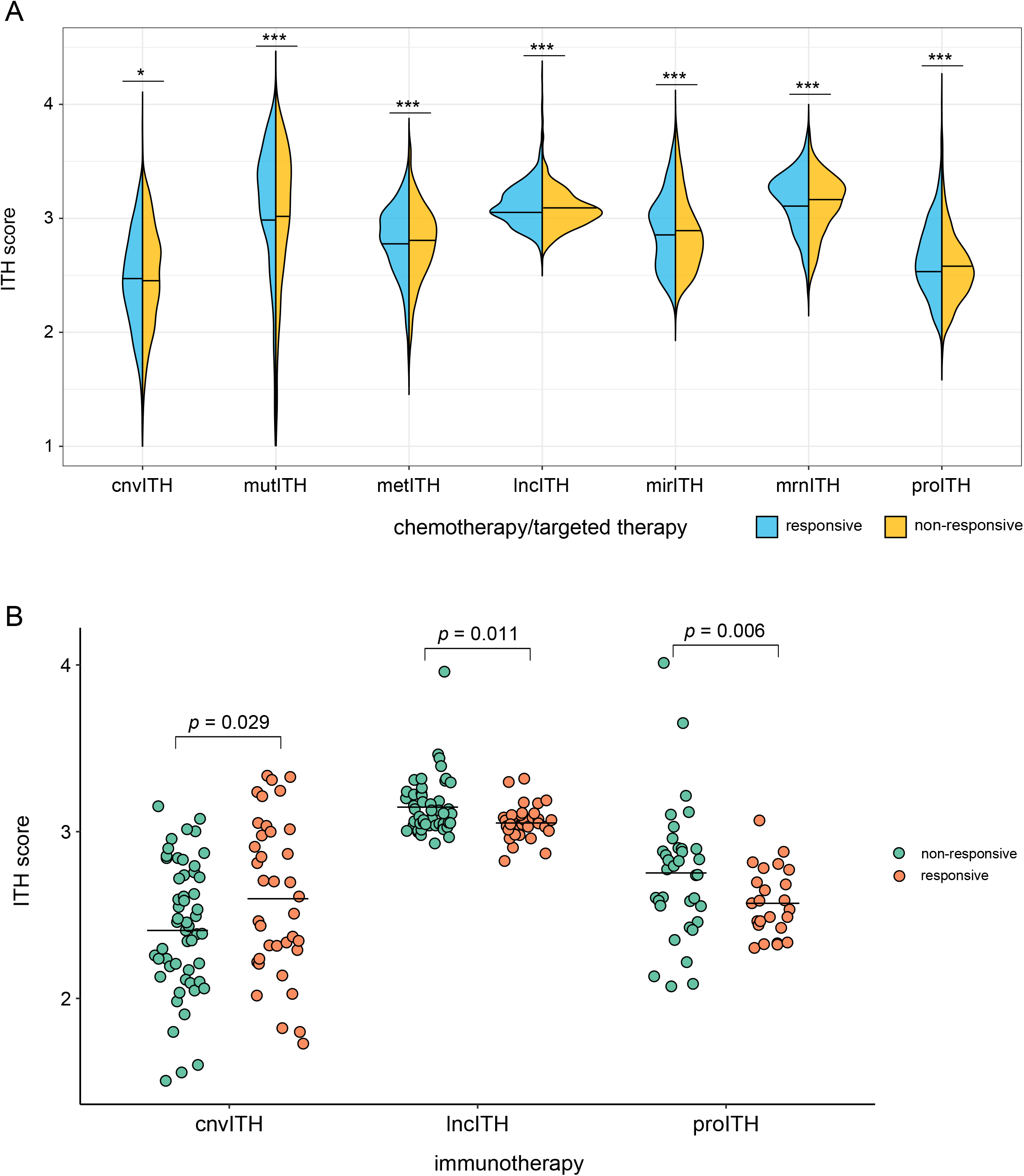
Correlations between the IE-based ITH scores and drug responses. **(A)** Six IE-based ITH scores are significantly lower in responsive than in non-responsive patients for chemotherapies or targeted therapies in pan-caner. **(B)** Two IE-based ITH scores are significantly lower in responsive than in non-responsive patients for immunotherapies in pancaner. The one-tailed Mann–Whitney *U* test *p* values are shown.

### The IE-based ITH likely correlates positively with tumor purity

We found that all the seven ITH scores had significant positive correlations with tumor purity in pan-cancer (Spearman correlation, *p* < 0.001) (Fig. 6A). In addition, seven and six ITH scores correlated positively with tumor purity in lung cancer and BRCA, respectively (*p* < 0.001) (Fig. 6A). These results suggest the rationality of our IE-based ITH evaluation algorithms since their ITH measures have captured the ITH within tumor cells.

**Fig. 6.**
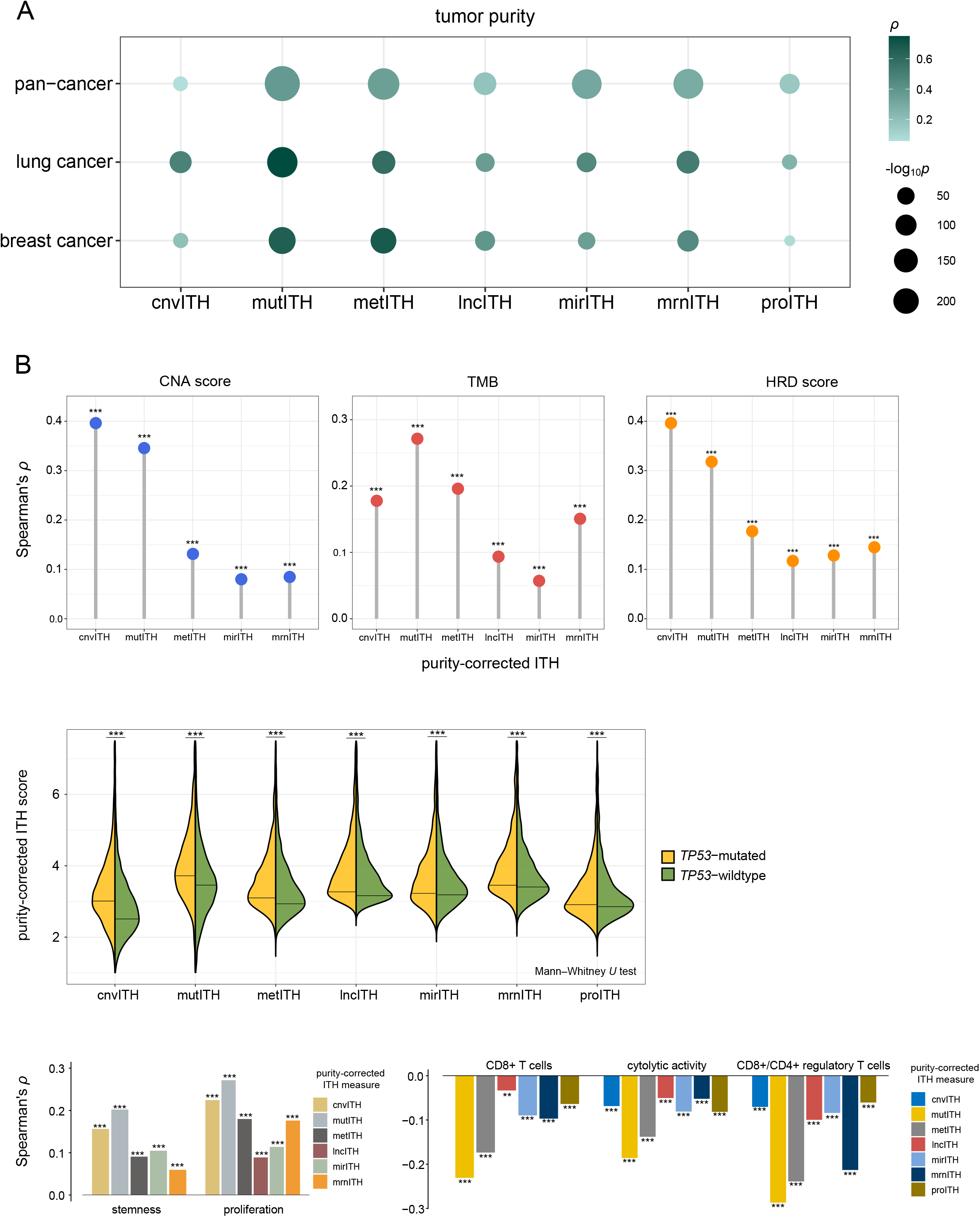
Correlations between the IE-based ITH scores and tumor purity. **(A)** The IE-based ITH scores correlate positively with tumor purity in pan-cancer, lung cancer, and breast cancer. **(B)** The tumor purity-corrected ITH scores correlate positively with genomic instability, tumor progression phenotypes, and antitumor immune responses. The Spearman correlation coefficients (*ρ*) and *p* values are shown.

To correct for the impact of tumor purity on the IE-based ITH scores in pan-cancer, we divided the ITH scores by tumor purity, which was the proportion of tumor cells in a bulk tumor and obtained from the TCGA pathological slides data. The results showed that the tumor purity-corrected ITH scores still had significant correlations with tumor progression, genomic instability, and antitumor immune responses (*p* < 0.01) (Fig. 6B).

### Associations among the IE-based ITH measures

We analyzed pairwise correlations among the seven IE-based ITH scores in pan-cancer and in the common cancer types. We found that most of these correlations were significant and positive (Spearman correlation, *p* < 0.05) (Fig. 7A and Supplementary Fig. S4). Among the 21 pairwise correlations, the correlation between mrnITH and lncITH scores was the strongest in pan-cancer (*ρ* = 0.66) and in BRCA (*ρ* = 0.86). These results are sensible since both mrnITH and lncITH measure ITH at the RNA level. In pan-cancer and in the common cancer types (except LGG), cnvITH scores had a stronger correlation with mutITH scores than with the other ITH scores (Fig. 7A and Supplementary Fig. S4). Again, it is justified since both cnvITH and mutITH measure ITH at the DNA level. metITH and mrnITH scores had the strongest correlation in GI cancer (*ρ* = 0.57) and kidney cancer ((*ρ* = 0.53) (Fig. 7A and Supplementary Fig. S4). Meanwhile, metITH scores had a stronger correlation with mutITH scores than with the other ITH scores in lung cancer (*ρ* = 0.70), GI cancer (*ρ* = 0.53), and BRCA (*ρ* = 0.64) (Fig. 7A and Supplementary Fig. S4). It is reasonable that metITH has strong correlations with both mrnITH and mutITH, because DNA methylation may regulate mRNA expression and meanwhile both metITH and mutITH measure ITH at the DNA level. Furthermore, because miRNA plays important roles in modulating mRNA expression, we anticipated that mirITH would have a strong correlation with mrnITH. Indeed, in pan-cancer and in the common cancer types (lung cancer, GI cancer, and BRCA), there were a relatively strong correlation between mirITH and mrnITH scores (*ρ* > 0.5).

**Fig. 7.**
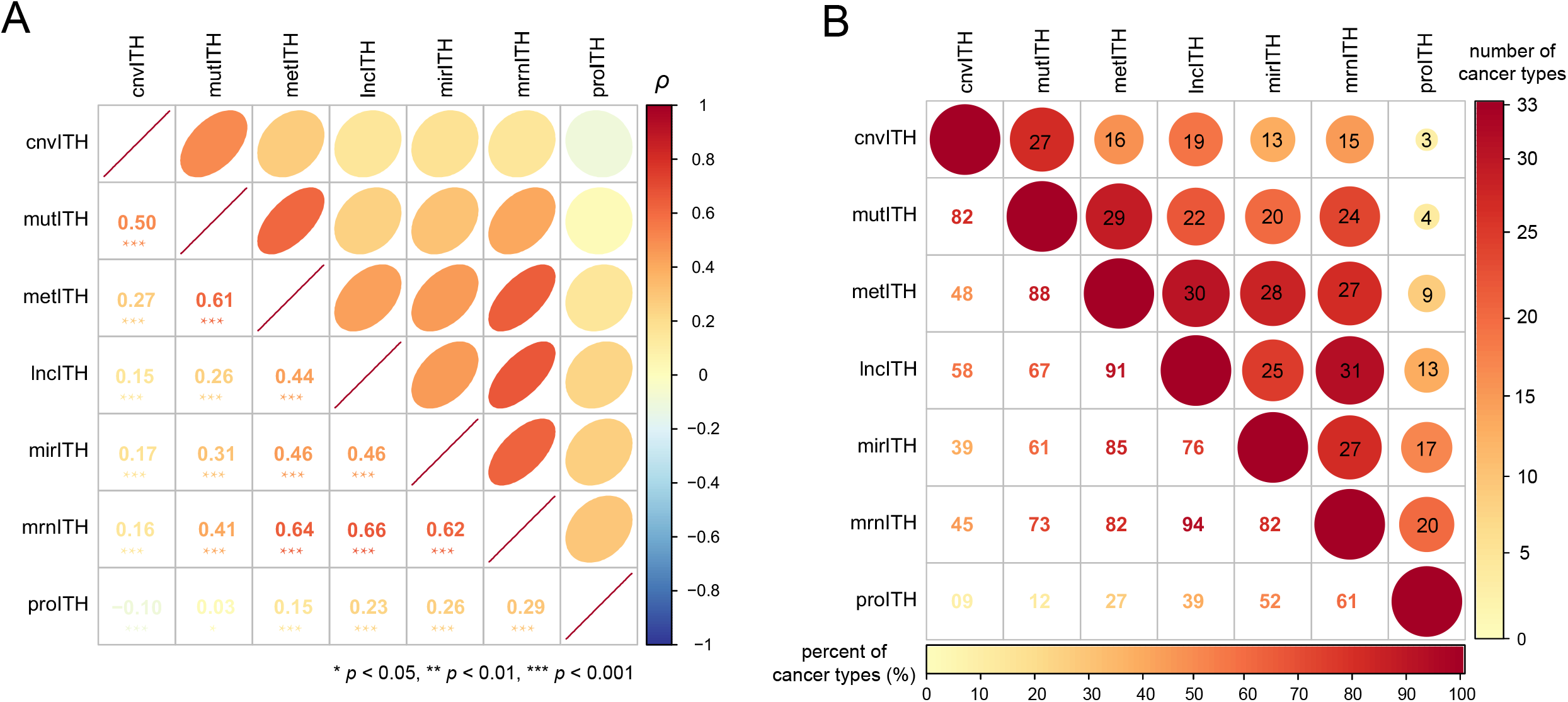
Pairwise correlations among the seven IE-based ITH scores. **(A)** Pairwise correlations among the seven IE-based ITH scores in pan-cancer. The Spearman correlation coefficients (*ρ*) and *p* values are shown. **(B)** Number of cancer types in which the pairwise correlations are significant among the 33 TCGA cancer types.

According to the central dogma of molecular biology proposed by Crick [23], we expected that the protein-level ITH would have a stronger correlation with the RNA-level ITH than with the DNA-level ITH. Indeed, proITH had a stronger correlation with mrnITH in pan-cancer (*ρ* = 0.29), lung cancer (*ρ* = 0.23), and kidney cancer (*ρ* = 0.34) than with the other ITH measures, and a stronger correlation with mirITH in GI cancer (*ρ* = 0.33), LGG (*ρ* = 0.45), and SKCM (*ρ* = 0.25) than with the other ITH measures (Fig. 7A and Supplementary Fig. S4). In contrast, proITH had the weakest correlation with cnvITH and mutITH.

We further analyzed pairwise correlations among the seven IE-based ITH scores in each of the 33 cancer types. We found that the correlation between mrnITH and lncITH scores was significant and positive in 31 cancer types (*p* < 0.05) (Fig. 7B). Besides, in at least 27 cancer types, there were significant positive correlations between lncITH and metITH, between mutITH and metITH, between metITH and mirITH, between mirITH and mrnITH, between metITH and mrnITH, and between cnvITH and mutITH. Overall, these results were in agreement with those by the prior analysis.

### Comparisons of the IE-based ITH measures across individual cancer types

We obtained the IE-based ITH scores for each of the 33 cancer types from pan-cancer and ranked them in order of their median ITH scores based on each of the seven ITH measure (Fig. 8A). In the 33 cancer types, THCA, THYM, DLBC, KICH and PCPG had the lowest median cnvITH scores, while UCS, OV, SARC, ESCA, and LUSC had the highest median cnvITH scores; THYM, PAAD, THCA, PCPG, TGCT, and LAML showed the lowest median mutITH scores, while UCS, READ, ESCA, SARC, and COAD showed the highest median mutITH scores. It indicates that THCA, THYM, and blood cancers (DLBC and LAML) have low ITH at the DNA level, as compared to gynecologic cancer (such as UCS and OV) and GI cancer (such as COAD, ESCA and READ) having high ITH at the DNA level. A direct explanation for these observations could be that gynecologic cancer and GI cancer are often with high genomic instability due to DNA repair deficiency caused by *BRCA* or *TP53* mutations [24-26], while THCA, THYM, and blood cancers have low genomic instability [27]. At the RNA level, PAAD, BRCA, THCA, PRAD, and KIRC likely had low ITH scores, while TGCT, LAML, UVM, DLBC, and ACC had high ITH scores. Similar results were also observed for DNA methylation and protein ITH. In general, THCA and PRAD were the cancer types with relatively low ITH scores across all the seven IE-based ITH measures. The low ITH could explain why these cancers generally have a relatively superior prognosis.

**Fig. 8.**
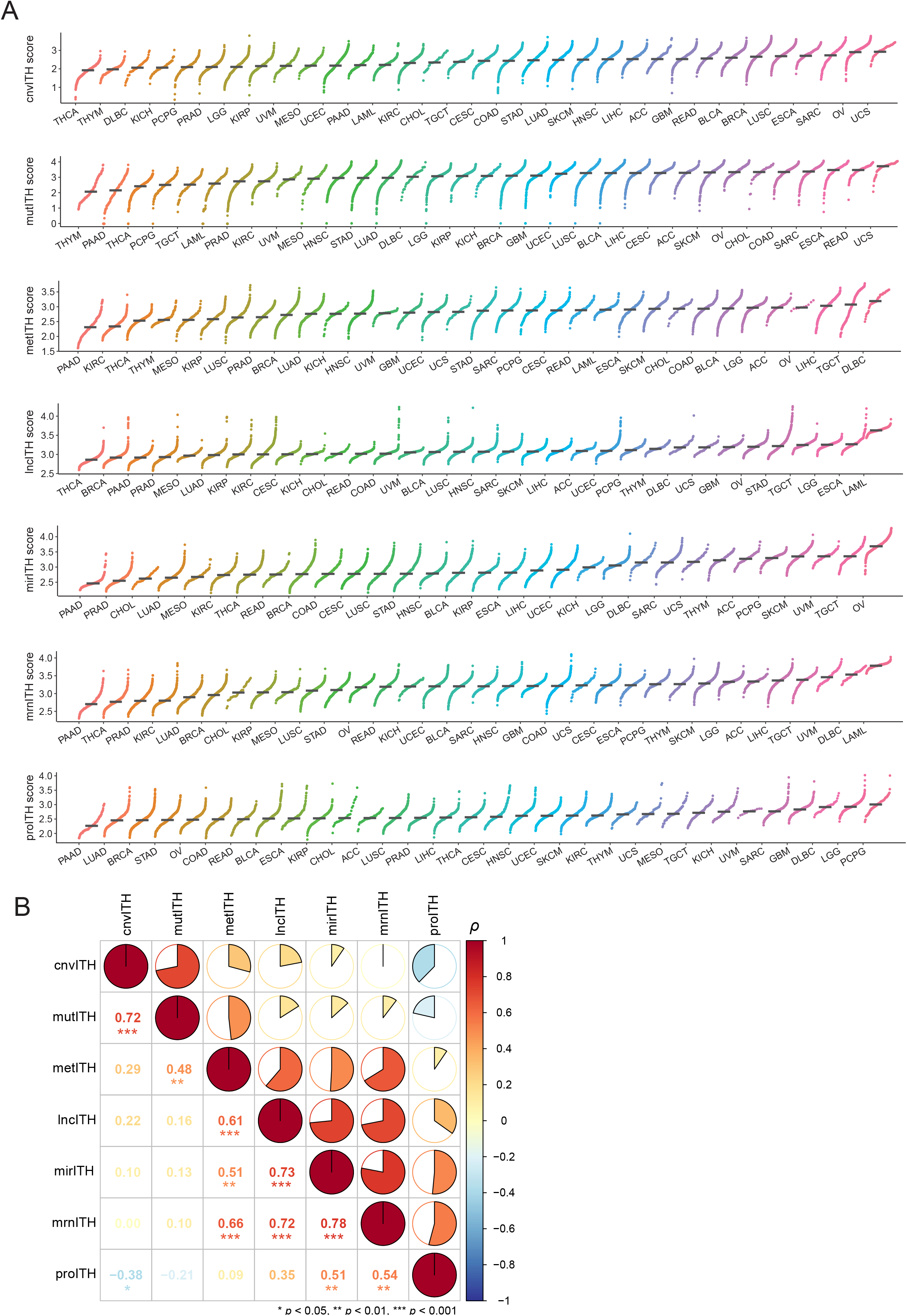
Comparisons of the IE-based ITH scores across individual cancer types. **(A)** The rank of 33 TCGA cancer types based on their median of ITH scores. **(B)** Pairwise correlations among the rank of 33 cancer types. The Spearman correlation coefficients (*ρ*) and *p* values are shown.

The median ITH score-based ranks generated a total of seven sets of ordinal numbers for the 33 cancer types. we evaluated correlations between pairwise sets of the ordinal numbers (Fig. 8B). For cnvITH, it had the strongest correlation with mutITH (*ρ* = 0.72; *p* < 0.001); metITH had the strongest correlation with mrnITH (*ρ* = 0.66; *p* < 0.001), which in turn had the strongest correlation with lncITH (*ρ* = 0.61; *p* < 0.001); lncITH had the strongest correlation with mirITH (*ρ* = 0.72; *p* < 0.001), which in turn had the strongest correlation with mrnITH (*ρ* = 0.61; *p* < 0.001); finally, proITH had the strongest correlation with mrnITH (*ρ* = 0.54) and mirITH (*ρ* = 0.51). Among all the correlations, the greatest two were 0.78 (between mrnITH and mirITH) and 0.72 (between mrnITH and lncITH). The direct regulatory relationships of miRNA and lncRNA with mrnITH may explain the greatest correlations between their ITH measures.

### Clustering analysis identifies four clusters of pan-cancer based on the IE-based ITH scores

Based on the seven types of IE-based ITH scores, two-way hierarchical clustering identified four clusters of pan-cancer, termed ITH-C1, ITH-C2, ITH-C3, and ITH-C4, respectively (Fig. 9A). ITH-C1 displayed the lowest scores of all the seven ITH measures; ITH-C2 showed relatively high scores of cnvITH and mutITH, but low scores of the other five ITH measures; ITH-C3 had high scores of all the seven ITH measures; ITH-C4 showed low scores of cnvITH and mutITH, but high scores of the other five ITH measures. Notably, survival analysis showed that ITH-C1 and ITH-C3 had the best and worst prognosis, respectively, for all the four endpoints (Fig. 9B). In addition, ITH-C4 likely had worse outcomes than ITH-C2 in PFI and DFI (*p* = 0.03 and 0.06, respectively). This analysis indicates that the RNA or protein-level ITH could be a stronger adverse prognosis factor than the DNA-level ITH.

**Fig. 9.**
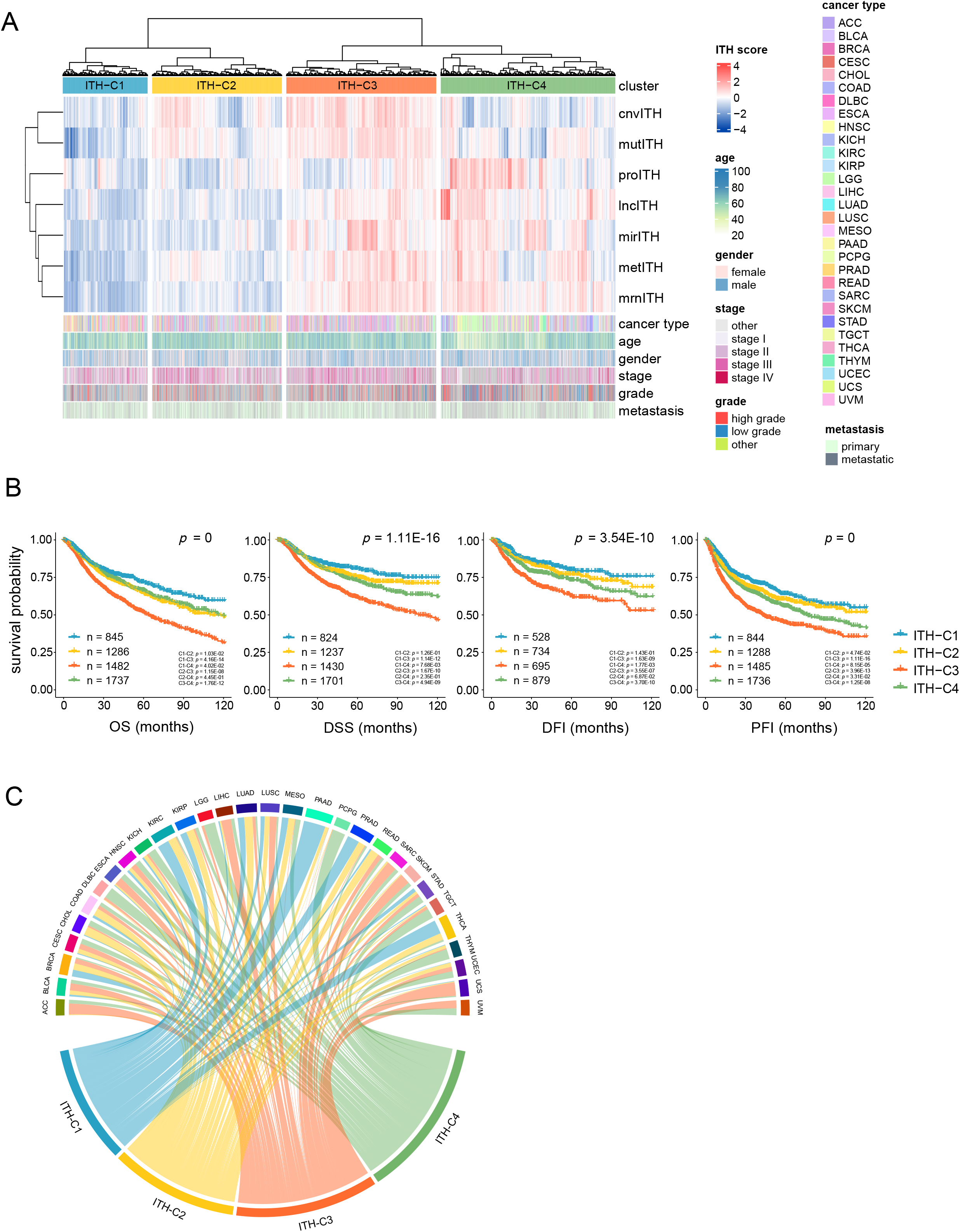
Two-way hierarchical clustering of tumor samples based on the seven types of IE-based ITH scores. **(A)** Clustering analysis identifies four clusters in pan-cancer: ITH-C1, ITH-C2, ITH-C3, and ITH-C4. **(B)** KM curves showing the significant different survival prognosis among the clustering subgroups. **(C)** The enrichment level of individual cancer types within a combination of cancer and cluster. Ro/e was used as the measure.

Furthermore, we compared distributions of 30 individual cancer types across the four clusters. Three cancer types (GBM, LAML, and OV) were excluded in this analysis because they had no any sample with all the seven ITH scores. We found that the sample distribution was significantly different for the 30 individual cancer types (chi-square test, *p* < 0.05). Next, we used the ratio of the numbers of observed to expected cancer samples in clusters to measure the enrichment of individual cancer types within the combination of cancer and cluster (Fig. 9C). We found that PAAD, THCA, PRAD, KIRC, and BRCA, had the highest proportions of samples in ITH-C1 (Ro/e = 4.94, 3.13, 2.72, 2.67, and 1.53, respectively), indicating that these cancers likely have the lowest ITH. Again, it conforms to the negative association between ITH and cancer prognosis since most of these cancers (except PAAD) had a relatively excellent prognosis. In ITH-C2, CHOL, COAD, READ, KIRP, and LUAD had the highest proportions of samples (Ro/e > 1.5). It indicates that these cancers have high ITH at the DNA level but low ITH at the RNA or protein level. In ITH-C3, ACC, BLCA, ESCA, LIHC, LUSC, SARC, SKCM, UCS, and UVM had the highest proportions of samples (Ro/e > 1.5), suggesting that these cancers have high ITH across all layers. Finally, in ITH-C4, DLBC, LGG, PCPG, TGCT, THYM, KICH, and UCEC had the highest proportions of samples (Ro/e > 1.5), suggesting their low ITH at the DNA level but high ITH at the RNA or protein level.

The two-way hierarchical clustering showed that cnvITH and mutITH belonged to a cluster (termed Cluster 1) and the other five ITH belonged to another cluster (termed Cluster 2). Further, Cluster 2 could be divided into two subclusters, with one subcluster including proITH and another subcluster including lncITH, mirITH, metITH, and mrnITH. These findings are consistent with previous ones obtained by analyzing pairwise correlations among the seven IE-based ITH scores in pan-cancer. Interestingly, mrnITH had the closest distance with metITH, supporting the important role of DNA methylation in regulating mRNA expression.

### Construction of an ITH measure by integrating the seven IE-based ITH measures

For each tumor sample, we built its ITH score by averaging its seven IE-based ITH scores after scaling into the range [0, 1] for each type of ITH measure. We termed the integrated ITH intITH. As expected, elevated intITH scores were correlated with worse clinical outcomes, tumor progression, genomic instability (increased CNA, TMB, and HRD, and *TP53* mutations), tumor immunosuppression (reduced enrichment of CD8+ T cells, cytolytic activity, and ratio of CD8+/CD4+ regulatory T cells), and higher tumor purity (Fig. 10A). To exhibit the excellence of intITH in characterizing ITH, we compared intITH with other ITH measures in their correlations with tumor progression phenotypes, antitumor immune response, clinical features, and genomic instability in the 33 TCGA cancer types. The other ITH measures included MATH [3], PhyloWGS [5], DITHER [6], DEPTH [2], and MYTH [8], which evaluate ITH based on profiles of somatic CNAs and/or mutations (MATH, DITHER, and PhyloWGS), mRNA expression (DEPTH), and DNA methylation (MYTH).

**Fig. 10.**
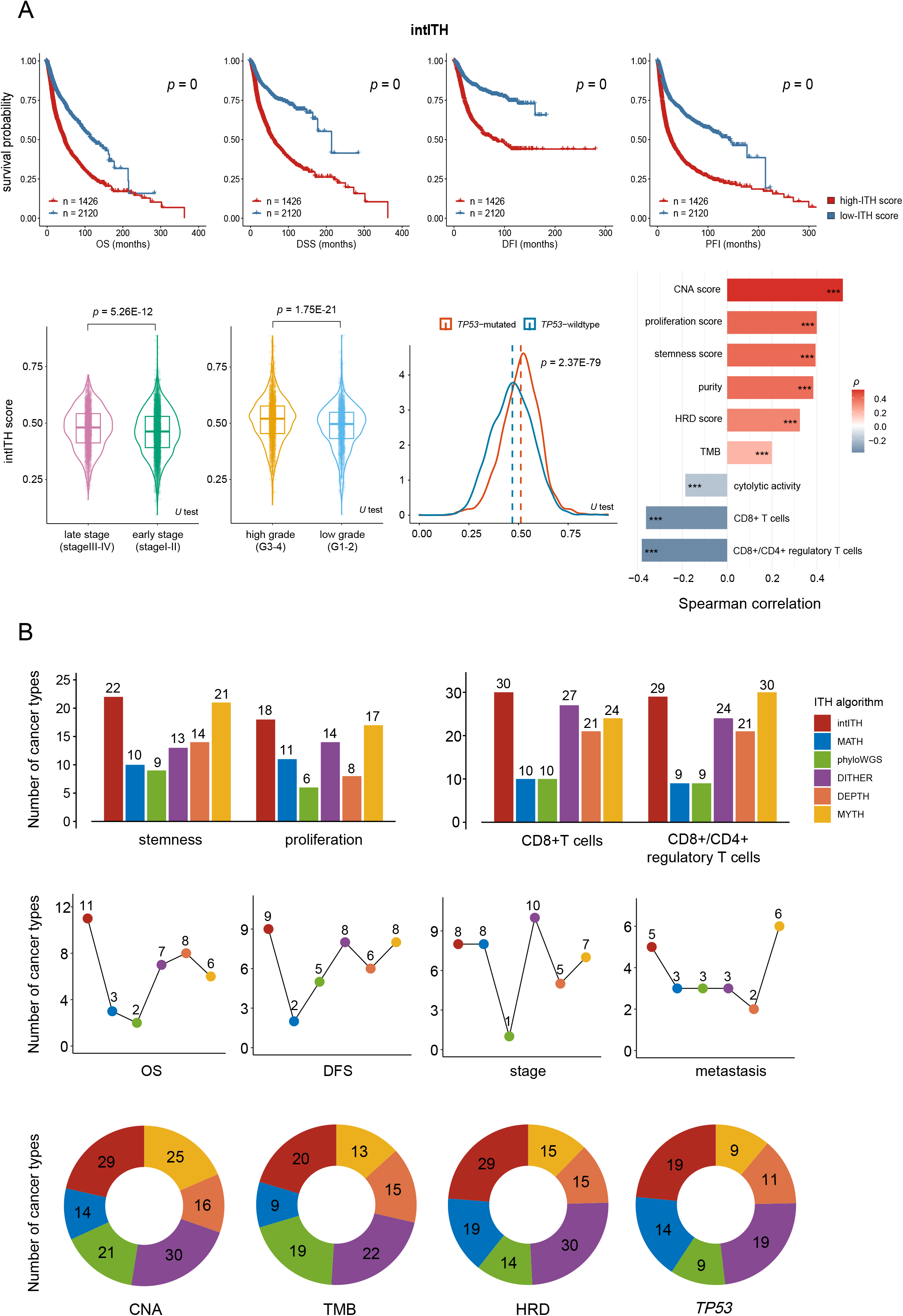
Integrating the seven IE-based ITH scores and comparing the integrated ITH with other five ITH evaluation algorithms. **(A)** The integrated ITH (intITH) showing significant correlations with unfavorable clinical outcomes, tumor progression, genomic instability (*TP53* mutations, high CNA scores, TMB, and HRD scores), tumor immunosuppression (lower enrichment of CD8+ T cells, cytolytic activity, and the ratio of CD8+/CD4+ regulatory T cells), and higher tumor purity. **(B)** Comparisons of intITH with five ITH evaluation algorithms (MATH, PhyloWGS, DITHER, DEPTH, and MYTH) in their correlations with tumor progression, immunosuppression, clinical outcomes, and genomic instability in 33 cancer types. The number of cancer types in which ITH scores have significant correlations with these features are given.

We found that stemness scores had significant positive correlations with ITH scores by MATH, PhyloWGS, DITHER, DEPTH, and MYTH in 10, 9, 13, 14, and 21 cancer types, respectively, compared to intITH in 22 cancer types (*p* < 0.05) (Fig. 10B). Proliferation scores had positive correlations with ITH scores by MATH, PhyloWGS, DITHER, DEPTH, and MYTH in 11, 6, 14, 8, and 17 cancer types, respectively, compared to intITH in 18 cancer types (*p* < 0.05) (Fig. 10B). These results suggest that the intITH measure has a stronger association with tumor progression phenotypes than the other ITH measures. The enrichment scores of CD8+ T cells had significant negative correlations with ITH scores by MATH, PhyloWGS, DITHER, DEPTH, and MYTH in 10, 10, 27, 21, and 24 cancer types, respectively, compared to intITH in 30 cancer types (*p* < 0.05) (Fig. 10B). Furthermore, intITH scores correlated inversely with the ratios of CD8+/CD4+ regulatory T cells in 29 cancer types, compared to MATH, PhyloWGS, DITHER, DEPTH, and in 9, 9, 24, 21, and 30 cancer types, respectively (*p* < 0.05) (Fig. 10B). These results indicate that antitumor immune responses are likely to have a stronger association with intITH than with the other algorithms’ ITH.

ITH scores by intITH were negatively correlated with OS time in 11 cancer types, compared to MATH, PhyloWGS, DITHER, DEPTH, and MYTH scores in 3, 2, 7, 8, and 6 cancer types, respectively (log-rank test, *p* < 0.05) (Fig. 10B). Moreover, intITH scores correlated inversely with disease-free survival (DFS) time in 9 cancer types, versus MATH, PhyloWGS, DITHER, DEPTH, and MYTH scores in 2, 5, 8, 6 and 8 cancer types, respectively (log-rank test, *p* < 0.05) (Fig. 10B). In 8 cancer types, intITH scores were significantly higher in late-stage than in early-stage tumors, versus MATH, PhyloWGS, DITHER, DEPTH, and MYTH scores in 8, 1, 10, 5 and 7 cancer types, respectively (*p* < 0.05) (Fig. 10B). Besides, in 5 cancer types, intITH scores were significantly higher in metastatic than in primary tumors, compared to MATH, PhyloWGS, DITHER, DEPTH, and MYTH scores in 3, 3, 3, 2 and 8 cancer types, respectively (*p* < 0.05) (Fig. 10B). Overall, these results suggest that intITH likely has a stronger association with unfavorable clinical outcomes in cancer than the other algorithms’ ITH.

We found that CNA scores had significant positive correlations with ITH scores by MATH, PhyloWGS, DITHER, DEPTH, and MYTH in 14, 21, 30, 16, and 25 cancer types, respectively, compared to intITH in 29 cancer types (*p* < 0.05) (Fig. 10B). TMB correlated positively with ITH scores by MATH, PhyloWGS, DITHER, DEPTH, and MYTH in 9, 19, 22, 15, and 13 cancer types, respectively, compared to intITH in 20 cancer types (*p* < 0.05) (Fig. 10B). Furthermore, HRD scores had significant positive correlations with ITH scores by MATH, PhyloWGS, DITHER, DEPTH, and MYTH in 19, 14, 30, 15, and 15 cancer types, respectively, compared to intITH in 29 cancer types (*p* < 0.05) (Fig. 10B). In addition, *TP53* mutations were associated with higher ITH scores by MATH, PhyloWGS, DITHER, DEPTH, and MYTH in 14, 9, 19, 11, and 9 cancer types, respectively, compared to intITH in 19 cancer types (*p* < 0.05) (Fig. 10B). Taken together, these results indicate that intITH has a stronger association with genomic instability than the other algorithms’ ITH except DITHER, which evaluates ITH based on both somatic mutation and CNA profiles.

Altogether, these results indicate that the intITH algorithm has more superior performance in characterizing ITH than the established algorithms.

## Discussion

Because ITH is associated with tumor progression, relapse, immunoevasion, and drug resistance, evaluation of ITH levels is meaningful in clinical practice. Existing algorithms for measuring ITH are limited to a single molecular level, e.g., DNA [3-6] and mRNA [2, 7]. In this study, we proposed a set of algorithms for measuring ITH at the DNA (genome), mRNA, miRNA, lncRNA, protein, and epigenome level, respectively. These algorithms were designed all based on an underlying concept, namely IE, which measures variables’ variation. We argue that the heterogeneity among tumor cells is not only demonstrated at the genome level, but also at the transcriptome, proteome, and even epigenome level. Indeed, all the IE-based ITH measures displayed the typical properties of ITH, such as their significant correlations with unfavorable clinical outcomes, tumor progression phenotypes, genomic instability, and antitumor immunosuppression. Furthermore, the integrated ITH measure by combining the seven ITH scores showed more prominent properties of ITH, as evidenced by its stronger associations with clinical outcomes, tumor progression, genomic features, and tumor immunity than the ITH measures at a single molecular level.

Correlation and clustering analysis consistently demonstrated that the correlations between ITH measures at identical molecular levels, such as cnvITH and mutITH, mrnITH and mirITH, and mrnITH and lncITH, were stronger than those at different molecular levels, such as cnvITH and proITH, and mutITH and proITH. Interestingly, the epigenome-based ITH (metITH) and mRNA-based ITH (mrnITH) had stronger correlations with mutITH than with cnvITH, although both mutITH and cnvITH measured ITH at the genome level. In addition, the mRNA-based ITH showed stronger correlations with mirITH, lncITH, and metITH than with mutITH and cnvITH in general, supporting the regulatory relationships of miRNA, lncRNA, and DNA methylation towards mRNA. Finally, the protein-level ITH (proITH) displayed stronger correlations with the RNA-level ITH (e.g., mrnITH) than with the DNA-level ITH, particularly the genome-level ITH. It supports the central dogma of molecular biology.

Our clustering analysis on the basis of the seven IE-based ITH measures revealed the heterogeneity of ITH among different cancer types. For example, nervous system tumors (such as LGG and PCPG) were predominant in ITH-C4, characterized by low genome-level ITH but high ITH at the RNA, protein, and epigenome levels. It conforms to the relatively low levels of genomic instability in nervous system tumors [27]. COAD and READ were predominant in ITH-C2, which contrasted with ITH-C4, characterized by high genome-level ITH but low ITH at the RNA, protein, and epigenome levels. Again, it is consistent with the high levels of genomic instability in colorectal cancer, which involves three types of genomic instability: microsatellite instability, chromosome instability, and chromosomal translocation [25]. Interestingly, the cancers originated from the same tissue or organ but different cells or parts may have significantly different ITH profiles. For example, two subtypes of lung cancer, LUAD and LUSC, were predominant in ITH-C2 and ITH-C3, respectively; three subtypes of kidney cancer, KIRC, KIRP, and KICH, were prevalent in ITH-C1, ITH-C2, and ITH-C4, respectively. Surprisingly, PAAD as a highly malignant tumor with an extremely inferior prognosis, was prevalent in ITH-C1, indicating the low ITH of this cancer across genome, transcriptome, proteome, and epigenome levels. However, PAAD is characterized by high TMB [27], with frequent somatic mutations in *KRAS* and *TP53* [28]. In contrast, THCA and PRAD are the cancer types with excellent prognosis, which were also predominant in ITH-C1, the subgroup with low ITH across all the molecular levels. Thus, the inverse association between ITH and prognosis is not perfectly established in some cases.

Here we regard each gene, mRNA, miRNA, lncRNA, protein, and methylation region (or CpG island) as the basic element of their respective molecular systems. Our analysis suggests that it is the variation of the molecular system’s perturbation rather than the molecular system’s perturbation itself that correlates with cancer development and outcomes. In fact, even though all or most elements display dramatic perturbations simultaneously to result in a large-scale perturbation of the molecular systems in cancer patients, they are likely to have favorable outcomes if the elements’ perturbations are homogeneous or harmonious to yield the low ITH.

This study is with several limitations. First, because the number of proteins is relatively small (< 300) in the TCGA datasets, the proITH scores may not fully reflect the level of ITH at the protein level. This could explain why the correlations of proITH scores with clinical outcomes, tumor progression, genomic features, and tumor immunity are generally less significant as compared to the other six ITH measures. Second, in most cases, we determined a significant correlation in the correlation analysis merely based on *p* values, but not take into account the correlation coefficient. Thus, the definition of strong, medium, and weak correlations through cut-offs of correlation coefficients would be more sensible. Third, because most bulk tumors are not composed of merely tumor cells, the ITH measures are likely confounded by non-tumor components. Thus, introduction of the variable of tumor purity into the algorithms could be more reasonable. Finally, we have not tested our algorithms using single-cell data, although multiomics data for cancer single cells remain scarce. Overcoming these limitations to improve our algorithms is the objective of our future investigation.

## Declarations

### Ethics approval and consent to participate

Not applicable.

### Consent for publication

Not applicable.

### Availability of data and materials

All data associated with this study are available within the paper and its supplementary data. The codes and scripts of the IE-based ITH algorithms (R package) are available at the website: https://github.com/WangX-Lab/ieITH.

## Acknowledgments

Not applicable.

## Author Contributions

HA performed the research, data analyses, and manuscript editing. DD performed the research, data analyses. XW conceived this study, designed the algorithm and analysis strategies, and wrote the manuscript. All the authors read and approved the final manuscript.

## List of Abbreviations

CN: copy number
CNA: copy number alteration
DDR: DNA damage repair
DFI: disease-free interval
DFS: disease-free survival
DSS: disease-specific survival
FA: Fanconi anemia
GDC: Genomic Data Commons
GI: gastrointestinal cancer
HRD: homologous recombination deficiency
ITH: intratumor heterogeneity
IE: Information entropy
KM: Kaplan–Meier
lncRNA: long non-coding RNA
MAF: mutant allele fraction
miRNA: microRNA
OS: overall survival
PFI: progression-free interval
ssGSEA: single-sample gene-set enrichment analysis
TCGA: The Cancer Genome Atlas
TMB: tumor mutation burden
ACC: adrenocortical carcinoma
BLCA: bladder urothelial carcinoma
BRCA: breast invasive carcinoma
CESC: cervical squamous cell carcinoma and endocervical adenocarcinoma
CHOL: cholangiocarcinoma
COAD: colon adenocarcinoma
DLBC: lymphoid neoplasm diffuse large B-cell lymphoma
ESCA: esophageal carcinoma
GBM: glioblastoma multiforme
HNSC: head and Neck squamous cell carcinoma
KICH: kidney chromophobe
KIRC: kidney renal clear cell carcinoma
KIRP: kidney renal papillary cell carcinoma
LAML: acute myeloid leukemia
LGG: brain lower grade glioma
LIHC: liver hepatocellular carcinoma
LUAD: lung adenocarcinoma
LUSC: lung squamous cell carcinoma
MESO: mesothelioma
OV: ovarian serous cystadenocarcinoma
PAAD: pancreatic adenocarcinoma
PCPG: pheochromocytoma and paraganglioma
PRAD: prostate adenocarcinoma
READ: rectum adenocarcinoma
SARC: sarcoma
SKCM: skin cutaneous melanoma
STAD: stomach adenocarcinoma
TGCT: testicular germ cell tumors
THCA: thyroid carcinoma
THYM: thymoma
UCEC: uterine corpus endometrial carcinoma
UCS: uterine carcinosarcoma
UVM: uveal melanoma.

