## Supplementary Information for "Defining Multiple Layers of Intratumor Heterogeneity Based on Variations of Perturbations in Multi-omics Profiling"

### Supplementary Data

**Supplementary Fig. S1 Correlations of the IE-based ITH measures with clinical and phenotypic features in the common cancer types. (A)** KM curves showing that patients with high-ITH scores (upper third) have better survival than those with low-ITH scores (bottom third) in the common cancer types. The log-rank test *p* values are shown. **(B)** IE-based ITH scores are significantly higher in late-stage (stage III-IV) than in early-stage (stage I-II) in lung cancer and in kidney cancer. **(C)** IE-based ITH scores are significantly higher in high-grade (G3-4) versus low-grade (G1-2) in glioma and in kidney cancer. **(D)** IE-based ITH scores are significantly higher in metastatic versus primary tumors in kidney cancer. The one-tailed Mann–Whitney *U* test *p* values are shown. **(E-F)** IE-based ITH scores are positively correlated with tumor stemness scores **(E)** and tumor cell proliferation scores in the common cancer types **(F)**. The Spearman correlation coefficients are shown. * $p < 0.05$, ** $p < 0.01$, *** $p < 0.001$.

**Supplementary Fig. S2 Correlations between the IE-based ITH scores and genomic instability in the common cancer types. (A)** IE-based ITH scores are positively correlated with CNA scores, TMB and HRD scores in the common cancer types. **(B)** IE-based ITH scores are positively correlated with DNA damage repair pathways in lung cancer. **(C)** Six types of ITH scores are significantly higher in *TP53*-mutated than in *TP53*-wildtype tumors in breast cancer, gastrointestinal cancer, and lung cancer. The one-tailed Mann–Whitney *U* test *p* values are shown.

**Supplementary Fig. S3 Correlations between the IE-based ITH scores and antitumor immune responses.** Most of the ITH scores are inversely correlated with the enrichment levels of CD8+ T cells, immune cytolytic activity and the ratios of CD8+/CD4+ regulatory T cells. Spearman correlation coefficients and *p* values are shown.

**Supplementary Fig. S4 Pairwise correlations among the seven IE-based ITH scores in the common cancer types.** Pairwise correlations among the seven IE-based ITH scores in the common cancer types. The Spearman correlation coefficients ($\rho$) and *p* values are shown.

**Supplementary Table S1 A summary of the multi-omics profiles used in this study.**

**Supplementary Table S2 The marker or pathway genes of immune signatures, biological processes, and pathways.**

### Supplementary Fig. S1

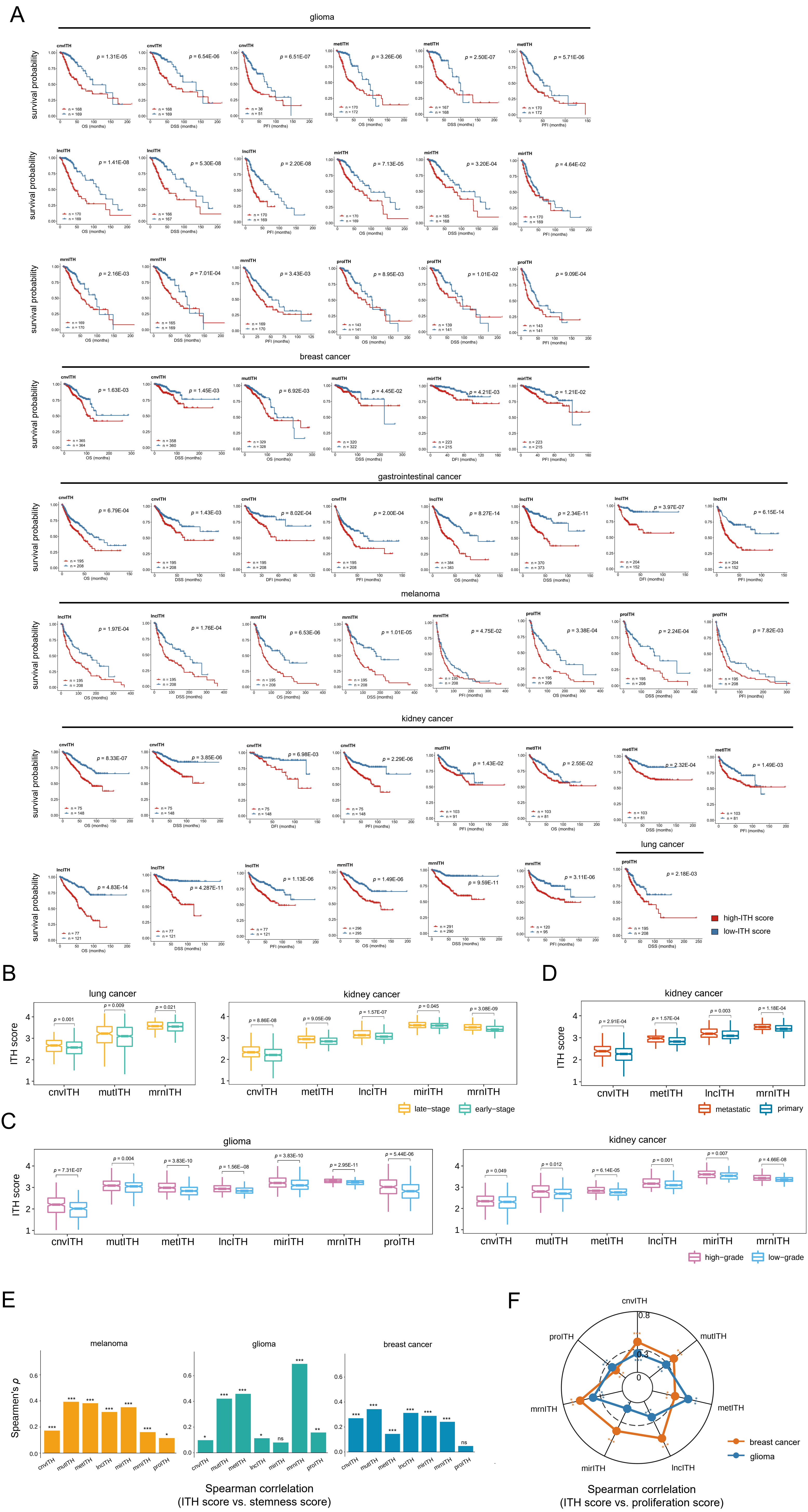

Supplementary Fig. S2

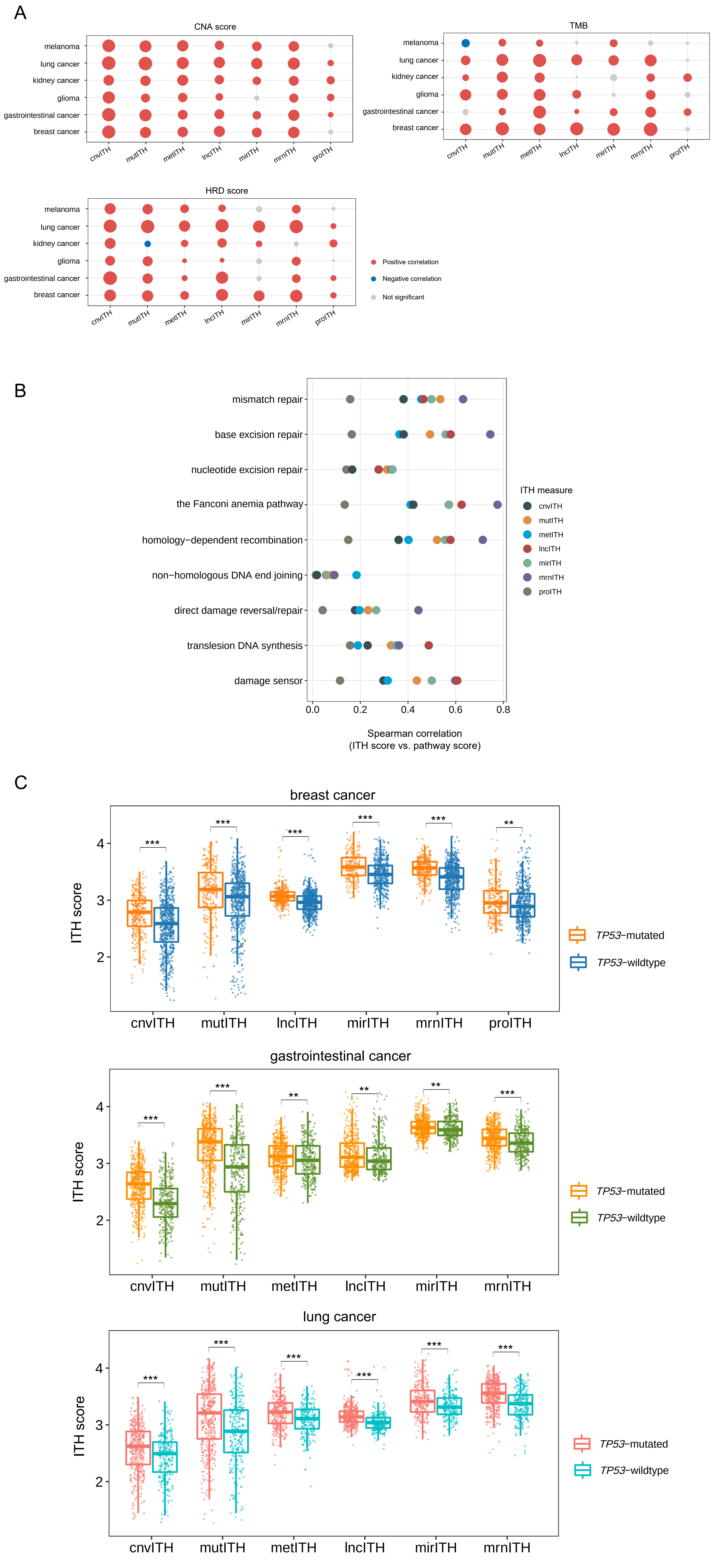

Supplementary Fig. S3

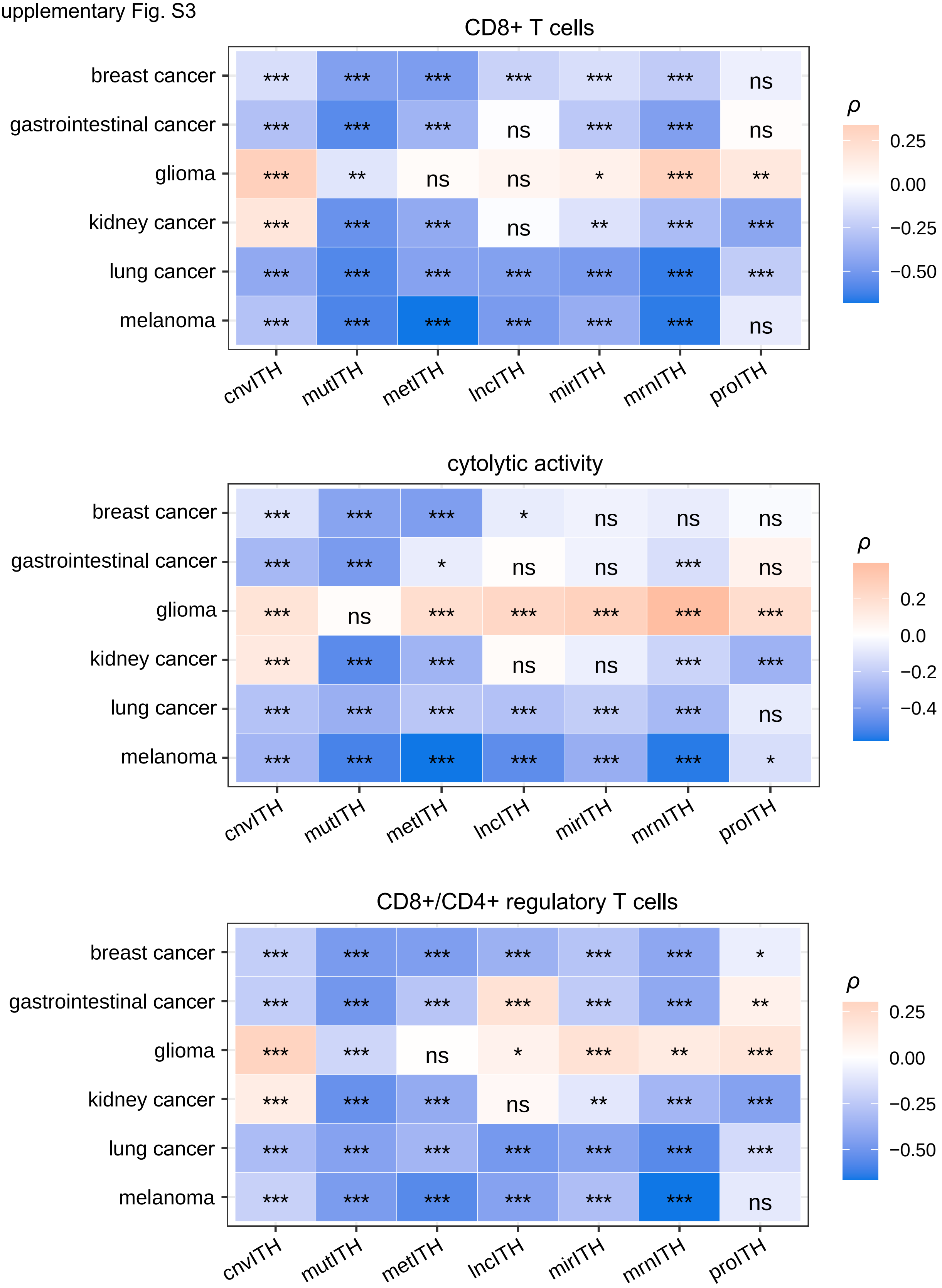

Supplementary Fig. S4

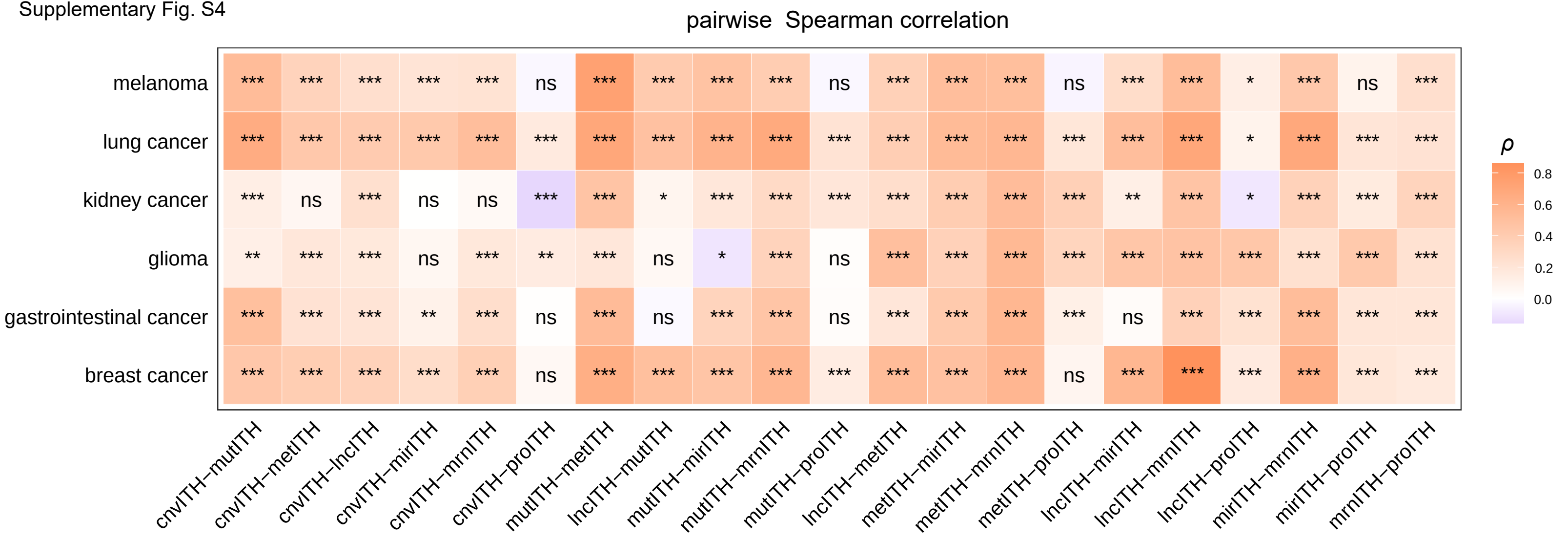

**Supplementary Table S1. A summary of the multi-omics profiles used in this study.**

| somatic mutations-associated "maf" files | | | | |
|---|---|---|---|---|
| **Cancer type** | **Dataset ID** | **Platform** | **Number of tumor samples** | **Sources** |
| Adrenocortical Carcinoma | TCGA-ACC | Illumina | 92 | TCGA |
| Bladder Urothelial Carcinoma | TCGA-BLCA | Illumina | 412 | TCGA |
| Breast Invasive Carcinoma | TCGA-BRCA | Illumina | 986 | TCGA |
| Cervical Squamous Cell Carcinoma and Endocervical Adenocarcinoma | TCGA-CESC | Illumina | 289 | TCGA |
| Cholangiocarcinoma | TCGA-CHOL | Illumina | 51 | TCGA |
| Colon Adenocarcinoma | TCGA-COAD | Illumina | 399 | TCGA |
| Lymphoid Neoplasm Diffuse Large B-cell Lymphoma | TCGA-DLBC | Illumina | 37 | TCGA |
| Esophageal Carcinoma | TCGA-ESCA | Illumina | 184 | TCGA |
| Glioblastoma Multiforme | TCGA-GBM | Illumina | 393 | TCGA |
| Head and Neck Squamous Cell Carcinoma | TCGA-HNSC | Illumina | 508 | TCGA |
| Kidney Chromophobe | TCGA-KICH | Illumina | 66 | TCGA |
| Kidney Renal Clear Cell Carcinoma | TCGA-KIRC | Illumina | 336 | TCGA |
| Kidney Renal Papillary Cell Carcinoma | TCGA-KIRP | Illumina | 281 | TCGA |
| Acute Myeloid Leukemia | TCGA-LAML | Illumina | 143 | TCGA |
| Brain Lower Grade Glioma | TCGA-LGG | Illumina | 508 | TCGA |
| Liver Hepatocellular Carcinoma | TCGA-LIHC | Illumina | 364 | TCGA |
| Lung Adenocarcinoma | TCGA-LUAD | Illumina | 567 | TCGA |
| Lung Squamous Cell Carcinoma | TCGA-LUSC | Illumina | 492 | TCGA |
| Mesothelioma | TCGA-MESO | Illumina | 82 | TCGA |
| Ovarian Serous Cystadenocarcinoma | TCGA-OV | Illumina | 436 | TCGA |
| Pancreatic Adenocarcinoma | TCGA-PAAD | Illumina | 178 | TCGA |
| Pheochromocytoma and Paraganglioma | TCGA-PCPG | Illumina | 179 | TCGA |
| Prostate Adenocarcinoma | TCGA-PRAD | Illumina | 495 | TCGA |
| Rectum Adenocarcinoma | TCGA-READ | Illumina | 137 | TCGA |
| Sarcoma | TCGA-SARC | Illumina | 237 | TCGA |
| Skin Cutaneous Melanoma | TCGA-SKCM | Illumina | 467 | TCGA |
| Stomach Adenocarcinoma | TCGA-STAD | Illumina | 437 | TCGA |
| Testicular Germ Cell Tumors | TCGA-TGCT | Illumina | 144 | TCGA |
| Thyroid Carcinoma | TCGA-THCA | Illumina | 492 | TCGA |
| Thymoma | TCGA-THYM | Illumina | 123 | TCGA |
| Uterine Corpus Endometrial Carcinoma | TCGA-UCEC | Illumina | 530 | TCGA |
| Uterine Carcinosarcoma | TCGA-UCS | Illumina | 57 | TCGA |
| Uveal Melanoma | TCGA-UVM | Illumina | 80 | TCGA |
| Pan-cancer | TCGA-PANCAN | Illumina | 10,182 | TCGA |

| CNAs-associated “SNP6” files | | | |
|---|---|---|---|
| **Dataset ID** | **Platform** | **Number of tumor samples** | **Sources** |
| TCGA-ACC | Affymetrix SNP 6.0 | 90 | TCGA |
| TCGA-BLCA | Affymetrix SNP 6.1 | 415 | TCGA |
| TCGA-BRCA | Affymetrix SNP 6.2 | 1,111 | TCGA |
| TCGA-CESC | Affymetrix SNP 6.3 | 297 | TCGA |
| TCGA-CHOL | Affymetrix SNP 6.4 | 36 | TCGA |
| TCGA-COAD | Affymetrix SNP 6.5 | 466 | TCGA |
| TCGA-DLBC | Affymetrix SNP 6.6 | 48 | TCGA |
| TCGA-ESCA | Affymetrix SNP 6.7 | 185 | TCGA |
| TCGA-GBM | Affymetrix SNP 6.8 | 613 | TCGA |
| TCGA-HNSC | Affymetrix SNP 6.9 | 524 | TCGA |
| TCGA-KICH | Affymetrix SNP 6.10 | 66 | TCGA |
| TCGA-KIRC | Affymetrix SNP 6.11 | 536 | TCGA |
| TCGA-KIRP | Affymetrix SNP 6.12 | 290 | TCGA |
| TCGA-LAML | Affymetrix SNP 6.13 | 200 | TCGA |
| TCGA-LGG | Affymetrix SNP 6.14 | 533 | TCGA |
| TCGA-LIHC | Affymetrix SNP 6.15 | 378 | TCGA |
| TCGA-LUAD | Affymetrix SNP 6.16 | 532 | TCGA |
| TCGA-LUSC | Affymetrix SNP 6.17 | 503 | TCGA |
| TCGA-MESO | Affymetrix SNP 6.18 | 87 | TCGA |
| TCGA-OV | Affymetrix SNP 6.19 | 606 | TCGA |
| TCGA-PAAD | Affymetrix SNP 6.20 | 185 | TCGA |
| TCGA-PCPG | Affymetrix SNP 6.21 | 183 | TCGA |
| TCGA-PRAD | Affymetrix SNP 6.22 | 502 | TCGA |
| TCGA-READ | Affymetrix SNP 6.23 | 166 | TCGA |
| TCGA-SARC | Affymetrix SNP 6.24 | 264 | TCGA |
| TCGA-SKCM | Affymetrix SNP 6.25 | 472 | TCGA |
| TCGA-STAD | Affymetrix SNP 6.26 | 442 | TCGA |
| TCGA-TGCT | Affymetrix SNP 6.27 | 156 | TCGA |
| TCGA-THCA | Affymetrix SNP 6.28 | 512 | TCGA |
| TCGA-THYM | Affymetrix SNP 6.29 | 124 | TCGA |
| TCGA-UCEC | Affymetrix SNP 6.30 | 545 | TCGA |
| TCGA-UCS | Affymetrix SNP 6.31 | 56 | TCGA |
| TCGA-UVM | Affymetrix SNP 6.32 | 80 | TCGA |
| TCGA-PANCAN | Affymetrix SNP 6.33 | 11,203 | TCGA |

| DNA methylation | | | | | | |
|---|---|---|---|---|---|---|
| **Dataset ID** | **Platform** | **Number of probes** | **Number of all samples** | **Number of tumor samples** | **Number of normal samples** | **Sources** |
| TCGA-ACC | Illumina Human Methylation 450 | 485,578 | 80 | 80 | 0 | TCGA |
| TCGA-BLCA | Illumina Human Methylation 450 | 485,578 | 434 | 413 | 21 | TCGA |
| TCGA-BRCA | Illumina Human Methylation 450 | 485,578 | 888 | 790 | 98 | TCGA |
| TCGA-CESC | Illumina Human Methylation 450 | 485,578 | 312 | 309 | 3 | TCGA |
| TCGA-CHOL | Illumina Human Methylation 450 | 485,578 | 45 | 36 | 9 | TCGA |
| TCGA-COAD | Illumina Human Methylation 450 | 485,578 | 337 | 299 | 38 | TCGA |
| TCGA-DLBC | Illumina Human Methylation 450 | 485,578 | 48 | 48 | 0 | TCGA |
| TCGA-ESCA | Illumina Human Methylation 450 | 485,578 | 202 | 186 | 16 | TCGA |
| TCGA-GBM | Illumina Human Methylation 450 | 485,578 | 155 | 153 | 2 | TCGA |
| TCGA-HNSC | Illumina Human Methylation 450 | 485,578 | 580 | 530 | 50 | TCGA |
| TCGA-KICH | Illumina Human Methylation 450 | 485,578 | 66 | 66 | 0 | TCGA |
| TCGA-KIRC | Illumina Human Methylation 450 | 485,578 | 480 | 320 | 160 | TCGA |
| TCGA-KIRP | Illumina Human Methylation 450 | 485,578 | 321 | 276 | 45 | TCGA |
| TCGA-LAML | Illumina Human Methylation 450 | 485,578 | 194 | 194 | 0 | TCGA |
| TCGA-LGG | Illumina Human Methylation 450 | 485,578 | 530 | 530 | 0 | TCGA |
| TCGA-LIHC | Illumina Human Methylation 450 | 485,578 | 429 | 379 | 50 | TCGA |
| TCGA-LUAD | Illumina Human Methylation 450 | 485,578 | 492 | 460 | 32 | TCGA |
| TCGA-LUSC | Illumina Human Methylation 450 | 485,578 | 415 | 372 | 43 | TCGA |
| TCGA-MESO | Illumina Human Methylation 450 | 485,578 | 87 | 87 | 0 | TCGA |
| TCGA-OV | Illumina Human Methylation 450 | 485,578 | 10 | 10 | 0 | TCGA |
| TCGA-PAAD | Illumina Human Methylation 450 | 485,578 | 195 | 185 | 10 | TCGA |
| TCGA-PCPG | Illumina Human Methylation 450 | 485,578 | 187 | 184 | 3 | TCGA |
| TCGA-PRAD | Illumina Human Methylation 450 | 485,578 | 549 | 499 | 50 | TCGA |
| TCGA-READ | Illumina Human Methylation 450 | 485,578 | 106 | 99 | 7 | TCGA |
| TCGA-SARC | Illumina Human Methylation 450 | 485,578 | 269 | 265 | 4 | TCGA |
| TCGA-SKCM | Illumina Human Methylation 450 | 485,578 | 476 | 474 | 2 | TCGA |
| TCGA-STAD | Illumina Human Methylation 450 | 485,578 | 398 | 396 | 2 | TCGA |
| TCGA-TGCT | Illumina Human Methylation 450 | 485,578 | 156 | 156 | 0 | TCGA |
| TCGA-THCA | Illumina Human Methylation 450 | 485,578 | 571 | 515 | 56 | TCGA |
| TCGA-THYM | Illumina Human Methylation 450 | 485,578 | 126 | 124 | 2 | TCGA |
| TCGA-UCEC | Illumina Human Methylation 450 | 485,578 | 478 | 432 | 46 | TCGA |
| TCGA-UCS | Illumina Human Methylation 450 | 485,578 | 57 | 57 | 0 | TCGA |
| TCGA-UVM | Illumina Human Methylation 450 | 485,578 | 80 | 80 | 0 | TCGA |
| TCGA-PANCAN | Illumina Human Methylation 450 | 485,578 | 9,736 | 8,990 | 746 | TCGA |

| mRNA expression | | | | | | |
|---|---|---|---|---|---|---|
| Dataset ID | Platform | Number of RNAs | Number of all samples | Number of tumor samples | Number of normal samples | Sources |
| TCGA-ACC | IlluminaHiSeq_RNASeqV2 | 20,531 | 205 | 77 | 128 | TCGA, GTEx |
| TCGA-BLCA | IlluminaHiSeq_RNASeqV2 | 20,531 | 427 | 408 | 19 | TCGA |
| TCGA-BRCA | IlluminaHiSeq_RNASeqV2 | 20,531 | 1,212 | 1,100 | 112 | TCGA |
| TCGA-CESC | IlluminaHiSeq_RNASeqV2 | 20,531 | 309 | 306 | 3 | TCGA |
| TCGA-CHOL | IlluminaHiSeq_RNASeqV2 | 20,531 | 45 | 36 | 9 | TCGA |
| TCGA-COAD | IlluminaHiSeq_RNASeqV2 | 20,531 | 328 | 287 | 41 | TCGA |
| TCGA-DLBC | IlluminaHiSeq_RNASeqV2 | 20,531 | 384 | 47 | 337 | TCGA, GTEx |
| TCGA-ESCA | IlluminaHiSeq_RNASeqV2 | 20,531 | 196 | 185 | 11 | TCGA |
| TCGA-GBM | IlluminaHiSeq_RNASeqV2 | 20,531 | 171 | 166 | 5 | TCGA |
| TCGA-HNSC | IlluminaHiSeq_RNASeqV2 | 20,531 | 566 | 522 | 44 | TCGA |
| TCGA-KICH | IlluminaHiSeq_RNASeqV2 | 20,531 | 91 | 66 | 25 | TCGA |
| TCGA-KIRC | IlluminaHiSeq_RNASeqV2 | 20,531 | 606 | 534 | 72 | TCGA |
| TCGA-KIRP | IlluminaHiSeq_RNASeqV2 | 20,531 | 323 | 291 | 32 | TCGA |
| TCGA-LAML | IlluminaHiSeq_RNASeqV2 | 20,531 | 173 | 173 | 0 | TCGA |
| TCGA-LGG | IlluminaHiSeq_RNASeqV2 | 20,531 | 1,664 | 523 | 1,141 | TCGA, GTEx |
| TCGA-LIHC | IlluminaHiSeq_RNASeqV2 | 20,531 | 423 | 373 | 50 | TCGA |
| TCGA-LUAD | IlluminaHiSeq_RNASeqV2 | 20,531 | 576 | 517 | 59 | TCGA |
| TCGA-LUSC | IlluminaHiSeq_RNASeqV2 | 20,531 | 552 | 501 | 51 | TCGA |
| TCGA-MESO | IlluminaHiSeq_RNASeqV2 | 20,531 | 87 | 87 | 0 | TCGA |
| TCGA-OV | IlluminaHiSeq_RNASeqV2 | 20,531 | 515 | 427 | 88 | TCGA, GTEx |
| TCGA-PAAD | IlluminaHiSeq_RNASeqV2 | 20,531 | 183 | 179 | 4 | TCGA |
| TCGA-PCPG | IlluminaHiSeq_RNASeqV2 | 20,531 | 187 | 184 | 3 | TCGA |
| TCGA-PRAD | IlluminaHiSeq_RNASeqV2 | 20,531 | 550 | 498 | 52 | TCGA |
| TCGA-READ | IlluminaHiSeq_RNASeqV2 | 20,531 | 105 | 95 | 10 | TCGA |
| TCGA-SARC | IlluminaHiSeq_RNASeqV2 | 20,531 | 265 | 263 | 2 | TCGA |
| TCGA-SKCM | IlluminaHiSeq_RNASeqV2 | 20,531 | 473 | 472 | 1 | TCGA |
| TCGA-STAD | IlluminaHiSeq_RNASeqV2 | 20,531 | 450 | 415 | 35 | TCGA |
| TCGA-TGCT | IlluminaHiSeq_RNASeqV2 | 20,531 | 319 | 154 | 165 | TCGA, GTEx |
| TCGA-THCA | IlluminaHiSeq_RNASeqV2 | 20,531 | 568 | 509 | 59 | TCGA |
| TCGA-THYM | IlluminaHiSeq_RNASeqV2 | 20,531 | 122 | 120 | 2 | TCGA |
| TCGA-UCEC | IlluminaHiSeq_RNASeqV2 | 20,531 | 381 | 370 | 11 | TCGA |
| TCGA-UCS | IlluminaHiSeq_RNASeqV2 | 20,531 | 135 | 57 | 78 | TCGA, GTEx |
| TCGA-UVM | IlluminaHiSeq_RNASeqV2 | 20,531 | 80 | 80 | 0 | TCGA |
| TCGA-PANCAN | IlluminaHiSeq_RNASeqV2 | 20,531 | 11,062 | 10,323 | 739 | TCGA |

| microRNA expression | | | | | | |
|---|---|---|---|---|---|---|
| **Dataset ID** | **Platform** | **Number of RNAs** | **Number of all samples** | **Number of tumor samples** | **Number of normal samples** | **Sources** |
| TCGA-ACC | IlluminaHiSeq_miRNASeq | 1,952 | 79 | 79 | 0 | TCGA |
| TCGA-BLCA | IlluminaHiSeq_miRNASeq | 2,210 | 429 | 410 | 19 | TCGA |
| TCGA-BRCA | IlluminaHiSeq_miRNASeq | 2,238 | 832 | 756 | 76 | TCGA |
| TCGA-CESC | IlluminaHiSeq_miRNASeq | 2,200 | 311 | 308 | 3 | TCGA |
| TCGA-CHOL | IlluminaHiSeq_miRNASeq | 1,779 | 45 | 36 | 9 | TCGA |
| TCGA-COAD | IlluminaHiSeq_miRNASeq | 2,113 | 261 | 253 | 8 | TCGA |
| TCGA-DLBC | IlluminaHiSeq_miRNASeq | 1,883 | 47 | 47 | 0 | TCGA |
| TCGA-ESCA | IlluminaHiSeq_miRNASeq | 2,092 | 195 | 183 | 12 | TCGA |
| TCGA-GBM | IlluminaHiSeq_miRNASeq | 0 | 0 | 0 | 0 | TCGA |
| TCGA-HNSC | IlluminaHiSeq_miRNASeq | 2,246 | 529 | 485 | 44 | TCGA |
| TCGA-KICH | IlluminaHiSeq_miRNASeq | 1,917 | 89 | 65 | 24 | TCGA |
| TCGA-KIRC | IlluminaHiSeq_miRNASeq | 2,048 | 311 | 241 | 70 | TCGA |
| TCGA-KIRP | IlluminaHiSeq_miRNASeq | 2,114 | 321 | 287 | 34 | TCGA |
| TCGA-LAML | IlluminaHiSeq_miRNASeq | 1,834 | 188 | 188 | 0 | TCGA |
| TCGA-LGG | IlluminaHiSeq_miRNASeq | 2,157 | 524 | 524 | 0 | TCGA |
| TCGA-LIHC | IlluminaHiSeq_miRNASeq | 2,172 | 420 | 371 | 49 | TCGA |
| TCGA-LUAD | IlluminaHiSeq_miRNASeq | 2,228 | 495 | 450 | 45 | TCGA |
| TCGA-LUSC | IlluminaHiSeq_miRNASeq | 2,213 | 380 | 336 | 44 | TCGA |
| TCGA-MESO | IlluminaHiSeq_miRNASeq | 1,964 | 87 | 87 | 0 | TCGA |
| TCGA-OV | IlluminaHiSeq_miRNASeq | 2,165 | 485 | 485 | 0 | TCGA |
| TCGA-PAAD | IlluminaHiSeq_miRNASeq | 2,050 | 182 | 178 | 4 | TCGA |
| TCGA-PCPG | IlluminaHiSeq_miRNASeq | 2,039 | 186 | 183 | 3 | TCGA |
| TCGA-PRAD | IlluminaHiSeq_miRNASeq | 2,111 | 544 | 492 | 52 | TCGA |
| TCGA-READ | IlluminaHiSeq_miRNASeq | 2,003 | 92 | 89 | 3 | TCGA |
| TCGA-SARC | IlluminaHiSeq_miRNASeq | 2,093 | 260 | 260 | 0 | TCGA |
| TCGA-SKCM | IlluminaHiSeq_miRNASeq | 2,220 | 452 | 450 | 2 | TCGA |
| TCGA-STAD | IlluminaHiSeq_miRNASeq | 2,178 | 428 | 387 | 41 | TCGA |
| TCGA-TGCT | IlluminaHiSeq_miRNASeq | 2,212 | 155 | 155 | 0 | TCGA |
| TCGA-THCA | IlluminaHiSeq_miRNASeq | 2,217 | 569 | 510 | 59 | TCGA |
| TCGA-THYM | IlluminaHiSeq_miRNASeq | 2,128 | 126 | 124 | 2 | TCGA |
| TCGA-UCEC | IlluminaHiSeq_miRNASeq | 2,238 | 430 | 399 | 31 | TCGA |
| TCGA-UCS | IlluminaHiSeq_miRNASeq | 2,010 | 56 | 56 | 0 | TCGA |
| TCGA-UVM | IlluminaHiSeq_miRNASeq | 1,938 | 80 | 80 | 0 | TCGA |
| TCGA-PANCAN | IlluminaHiSeq_miRNASeq | 2,454 | 9,405 | 8,766 | 639 | TCGA |

| long non-coding RNA expression | | | | | | |
|---|---|---|---|---|---|---|
| **Dataset ID** | **Platform** | **Number of RNAs** | **Number of all samples** | **Number of tumor samples** | **Number of normal samples** | **Sources** |
| TCGA-ACC | RNA-Seq | 14,805 | 79 | 79 | 0 | TCGA |
| TCGA-BLCA | RNA-Seq | 14,805 | 430 | 411 | 19 | TCGA |
| TCGA-BRCA | RNA-Seq | 14,805 | 1,217 | 1,104 | 113 | TCGA |
| TCGA-CESC | RNA-Seq | 14,805 | 309 | 306 | 3 | TCGA |
| TCGA-CHOL | RNA-Seq | 14,805 | 45 | 36 | 9 | TCGA |
| TCGA-COAD | RNA-Seq | 14,805 | 512 | 471 | 41 | TCGA |
| TCGA-DLBC | RNA-Seq | 14,805 | 48 | 48 | 0 | TCGA |
| TCGA-ESCA | RNA-Seq | 14,805 | 173 | 162 | 11 | TCGA |
| TCGA-GBM | RNA-Seq | 14,805 | 173 | 168 | 5 | TCGA |
| TCGA-HNSC | RNA-Seq | 14,805 | 546 | 502 | 44 | TCGA |
| TCGA-KICH | RNA-Seq | 14,805 | 89 | 65 | 24 | TCGA |
| TCGA-KIRC | RNA-Seq | 14,805 | 607 | 535 | 72 | TCGA |
| TCGA-KIRP | RNA-Seq | 14,805 | 321 | 289 | 32 | TCGA |
| TCGA-LAML | RNA-Seq | 14,805 | 151 | 151 | 0 | TCGA |
| TCGA-LGG | RNA-Seq | 14,805 | 529 | 529 | 0 | TCGA |
| TCGA-LIHC | RNA-Seq | 14,805 | 424 | 374 | 50 | TCGA |
| TCGA-LUAD | RNA-Seq | 14,805 | 585 | 526 | 59 | TCGA |
| TCGA-LUSC | RNA-Seq | 14,805 | 550 | 501 | 49 | TCGA |
| TCGA-MESO | RNA-Seq | 14,805 | 86 | 86 | 0 | TCGA |
| TCGA-OV | RNA-Seq | 14,805 | 379 | 379 | 0 | TCGA |
| TCGA-PAAD | RNA-Seq | 14,805 | 182 | 178 | 4 | TCGA |
| TCGA-PCPG | RNA-Seq | 14,805 | 186 | 183 | 3 | TCGA |
| TCGA-PRAD | RNA-Seq | 14,805 | 551 | 499 | 52 | TCGA |
| TCGA-READ | RNA-Seq | 14,805 | 177 | 167 | 10 | TCGA |
| TCGA-SARC | RNA-Seq | 14,805 | 265 | 263 | 2 | TCGA |
| TCGA-SKCM | RNA-Seq | 14,805 | 472 | 471 | 1 | TCGA |
| TCGA-STAD | RNA-Seq | 14,805 | 407 | 375 | 32 | TCGA |
| TCGA-TGCT | RNA-Seq | 14,805 | 156 | 156 | 0 | TCGA |
| TCGA-THCA | RNA-Seq | 14,805 | 568 | 510 | 58 | TCGA |
| TCGA-THYM | RNA-Seq | 14,805 | 121 | 119 | 2 | TCGA |
| TCGA-UCEC | RNA-Seq | 14,805 | 583 | 548 | 35 | TCGA |
| TCGA-UCS | RNA-Seq | 14,805 | 56 | 56 | 0 | TCGA |
| TCGA-UVM | RNA-Seq | 14,805 | 80 | 80 | 0 | TCGA |
| TCGA-PANCAN | RNA-Seq | 14,805 | 14,805 | 10,277 | 730 | TCGA |

| Protein expression | | | | | | |
|---|---|---|---|---|---|---|
| Dataset ID | Platform | Number of proteins | Number of all samples | Number of tumor samples | Number of normal samples | Sources |
| TCGA-ACC | RPPA | 192 | 46 | 46 | 0 | TCGA |
| TCGA-BLCA | RPPA | 208 | 344 | 344 | 0 | TCGA |
| TCGA-BRCA | RPPA | 226 | 898 | 898 | 0 | TCGA |
| TCGA-CESC | RPPA | 192 | 173 | 173 | 0 | TCGA |
| TCGA-CHOL | RPPA | 192 | 30 | 30 | 0 | TCGA |
| TCGA-COAD | RPPA | 208 | 362 | 362 | 0 | TCGA |
| TCGA-DLBC | RPPA | 192 | 33 | 33 | 0 | TCGA |
| TCGA-ESCA | RPPA | 192 | 126 | 126 | 0 | TCGA |
| TCGA-GBM | RPPA | 208 | 244 | 244 | 0 | TCGA |
| TCGA-HNSC | RPPA | 160 | 212 | 212 | 0 | TCGA |
| TCGA-KICH | RPPA | 193 | 63 | 63 | 0 | TCGA |
| TCGA-KIRC | RPPA | 217 | 478 | 478 | 0 | TCGA |
| TCGA-KIRP | RPPA | 195 | 216 | 216 | 0 | TCGA |
| TCGA-LAML | RPPA | 0 | 0 | 0 | 0 | TCGA |
| TCGA-LGG | RPPA | 201 | 435 | 435 | 0 | TCGA |
| TCGA-LIHC | RPPA | 219 | 184 | 184 | 0 | TCGA |
| TCGA-LUAD | RPPA | 223 | 365 | 365 | 0 | TCGA |
| TCGA-LUSC | RPPA | 223 | 328 | 328 | 0 | TCGA |
| TCGA-MESO | RPPA | 193 | 63 | 63 | 0 | TCGA |
| TCGA-OV | RPPA | 208 | 436 | 436 | 0 | TCGA |
| TCGA-PAAD | RPPA | 195 | 123 | 123 | 0 | TCGA |
| TCGA-PCPG | RPPA | 192 | 82 | 82 | 0 | TCGA |
| TCGA-PRAD | RPPA | 195 | 352 | 352 | 0 | TCGA |
| TCGA-READ | RPPA | 208 | 131 | 131 | 0 | TCGA |
| TCGA-SARC | RPPA | 192 | 226 | 226 | 0 | TCGA |
| TCGA-SKCM | RPPA | 208 | 355 | 355 | 0 | TCGA |
| TCGA-STAD | RPPA | 195 | 357 | 357 | 0 | TCGA |
| TCGA-TGCT | RPPA | 192 | 122 | 122 | 0 | TCGA |
| TCGA-THCA | RPPA | 175 | 224 | 224 | 0 | TCGA |
| TCGA-THYM | RPPA | 192 | 90 | 90 | 0 | TCGA |
| TCGA-UCEC | RPPA | 208 | 440 | 440 | 0 | TCGA |
| TCGA-UCS | RPPA | 192 | 48 | 48 | 0 | TCGA |
| TCGA-UVM | RPPA | 192 | 12 | 12 | 0 | TCGA |
| TCGA-PANCAN | RPPA | 258 | 7,744 | 7,744 | 0 | TCGA |

**Supplementary Table S2. The marker or pathway genes of immune signatures, biological processes, and pathways.**

| Immune signatures/ biological processes/ pathways | Gene set |
|---|---|
| CD8+ T cells | *CD2, CD247, CD28, CD3D, CD3E, CD3G, CD8A, ICAM1, ITGAL, ITGB2, PTPRC, THY1* |
| Cytolytic activity | *PRF1, GZMA* |
| CD4+ regulatory T cells | *CTLA4, FOXP3, GPR15, IL32, IL4, IL5* |
| Stemness | *DNMT3B, PFAS, XRCC5, HAUS6, TET1, IGF2BP1, PLAA, TEX10, MSH6, DLGAP5, SKIV2L2, SOHLH2, RRAS2, PAICS, CPSF3, LIN28B, IPO5, BMPR1A, ZNF788, ASCC3, FANCB, HMGA2, TRIM24, ORC1, HDAC2, HESX1, INHBE, MIS18A, DCUN1D5, MRPL3, CENPH, MYCN, HAUS1, GDF3, TBCE, RIOK2, BCKDHB,, RAD1, NREP, ADH5, PLRG1, ROR1, RAB3B, DIAPH3, GNL2, FGF2, NMNAT2, KIF20A, CENPI, DDX1, XXYLT1, GPR176, BBS9, C14orf166, BOD1, CDC123, SNRPD3, FAM118B, DPH3, EIF2B3, RPF2, APLP1, DACT1, PDHB, C14orf119, DTD1, SAMM50, CCL26, MED20, UTP6, RARS2, ARMCX2, RARS, MTHFD2, DHX15, HTR7, MTHFD1L, ARMC9, XPOT, IARS, HDX, ACTRT3, ERCC2, TBC1D16, GARS, KIF7, UBE2K, SLC25A3, ICMT, UGGT2, ATP11C, SLC24A1, EIF2AK4, GPX8, ALX1, OSTC, TRPC4, HAS2, FZD2, TRNT1, MMADHC, SNX8, CDH6, HAT1, SEC11A, DIMT1, TM2D2, FST, GBE1* |
| Proliferation | *CCNB1, CDC20, CDKN3, CDK1, MAD2L1, PRC1, RRM2* |
| Base excision repair pathway | *PARP1, POLB, APEX1, APEX2, FEN1, TDG, TDP1, UNG* |
| Nucleotide excision repair pathway | *CUL5, ERCC1, ERCC2, ERCC4, ERCC5, ERCC6, POLE, POLE3, XPA, XPC* |
| Mismatch repair pathway | *EXO1, MLH1, MLH3, MSH2, MSH3, MSH6, PMS1, PMS2* |
| the Fanconi anemia pathway | *FANCA, FANCB, FANCC, FANCD2, FANCI, FANCL, FANCM, UBE2T* |
| Homologous recombination pathway | *MRE11A, NBN, RAD50, TP53BP1, XRCC2, XRCC3, BARD1, BLM, BRCA1, BRCA2, BRIP1, EME1, GEN1, MUS81, PALB2, RAD51, RAD52, RBBP8, SHFM1, SLX1A, TOP3A* |
| Non-homologous end joining pathway | *LIG4, NHEJ1, POLL, POLM, PRKDC, XRCC4, XRCC5, XRCC6* |
| Direct repair pathway | *ALKBH2, ALKBH3, MGMT* |
| Translesion synthesis pathway | *POLN, POLQ, REV1, REV3L, SHPRH* |
| Damage sensor pathway | *ATM, ATR, ATRIP, CHEK1, CHEK2, MDC1, RNMT, TOPBP1, TREX1* |
